# Ether phospholipids modulate somatosensory responses by tuning multiple receptor functions in *Drosophila*

**DOI:** 10.1101/2023.09.12.556286

**Authors:** Takuto Suito, Kohjiro Nagao, Xiangmei Deng, Shoma Sato, Christian Ganser, Takayuki Uchihashi, Motosuke Tsutsumi, Tomomi Nemoto, Yuji Hara, Makoto Tominaga, Takaaki Sokabe

## Abstract

Transient receptor potential (TRP) and PIEZO channels are known receptors for physical stimuli, such as temperature and mechanical touch, respectively, in sensory nerves. As these receptors are localized in the plasma membrane, the modulation of sensory receptor activity by plasma membrane lipids has recently attracted attention. In this study, we focused on ether phospholipids (ePLs), which are abundant in neurons, and analyzed their role in somatosensation using *Drosophila* as a model. Reduced mechanosensory behavior was observed with ePL synthesizing gene knockout or knockdown in mechano-sensitive PIEZO-expressing neurons. The activation of PIEZO channels was significantly augmented in the presence of ePLs. Furthermore, we observed that ePLs modulate the thermosensory behavior and reduce thermal threshold of the thermosensitive TRPA1 channels. Finally, we revealed that ePLs affect membrane tension and lipid order of the plasma membrane in culture cells. Our study identified ePLs as a modulator of multiple somatosensation modalities in *Drosophila*, which underscore the significance of functional interaction between membrane lipid and sensory channel proteins.

## INTRODUCTION

The sensory system is indispensable to detect environmental parameters. Sensory perception is initiated by the activation of sensory receptors in several types of cells, including sensory nerves, which convert external stimuli into electrical or chemical signals. Transient receptor potential (TRP) channels^1–3^ and PIEZO channels^4^ are considered receptors for physical stimuli, such as temperature and mechanical touch, respectively, and their roles in sensory processing and the underlying mechanisms have been extensively studied^5–7^.

As these receptors are membrane proteins, the functional interaction between ion channels and membrane lipids has been key for resolving their activation mechanisms and cellular functions^8^. PIEZO channel activities are modulated by a variety of lipids. The activity of PIEZO1 is affected by the enrichment of either saturated or polyunsaturated FA (PUFAs) in membrane phospholipids^9,10^. Phosphatidylserine (PS) may interact with PIEZO1, and the asymmetric distribution of PS in the lipid bilayer is critical for the channel activity^11^. Similarly, the activity of many TRP channels is regulated by a wide variety of lipid molecules, such as phosphatidylinositol phosphates, fatty acids (FA) and their metabolites, endocannabinoids, lysophospholipids, and sphingolipids^12–15^. The modes of action of these lipids are either direct or indirect, modulating PIEZO and TRP channel activity positively or negatively depending on cellular context and signaling pathways. These tight functional linkages between sensory ion channels and lipid molecules exemplify their importance in cellular functions.

Despite many reports on the interaction between sensory receptors and lipids at a cellular level, a limited number of studies evaluated the physiological significance of lipid molecules on sensory responses in animals. For example, PUFA-containing phospholipids are highly enriched and play a significant role in the brain^16,17^. In *C. elegans*, arachidonic acid-containing phospholipids modulate the function of touch receptor neurons^18^. In *Drosophila*, an increase in the content of linoleic acid in sensory neurons changes TRPA1 activity and the preferred temperature of larvae^19^. Recently, it has been reported that linoleic acid increases PIEZO2 activity, and that linoleic acid supplementation improves Angelman Syndrome-associated mechanosensory deficits in mice^10^. These results suggest that lipid molecules may modulate the responsiveness and sensitivity of ion channels, thereby influencing neural function and the consequent physiological outputs.

An interesting lipid species in neural tissues is an ether phospholipid (ePL) identified in cancer cells in the 1960s^20^. ePLs comprise 20% of the lipid content in the mouse central nervous system (CNS), appearing as plasmalogen-type phosphatidylethanolamine (PE)^21^. In addition, ePLs have been implicated in neurodegenerative diseases; for instance, lower plasmalogen ePL content has been reported in Alzheimer’s disease, Parkinson’s disease, and Huntington’s disease^22^. Although the causal relationship between disease progression and the decrease in ePLs has not been clarified, they may be important for neural function. Physicochemical and molecular biological analyses indicated that ePL alters cell membrane properties^23,24^ and channel protein activity^25^. However, so far, the physiological significance of these processes has not been clearly elucidated. Furthermore, it is unclear whether ePLs are functionally involved in peripheral neurons.

Here, we combined lipidomic, behavioral, electrophysiological analyses, and measurements of the physicochemical properties of cell membranes to demonstrate the functional linkage between ePLs and membrane proteins in *Drosophila* sensory functions. Lipid analysis revealed the existence of ePLs, particularly ether phosphatidylethanolamines (ePEs), in neural tissues. Then, we analyzed somatosensory behaviors in larvae and observed that a knockout or sensory neuron-specific knockdown of alkylglycerone phosphate synthase (*AGPS*), an essential enzyme for ePL biosynthesis^26^, decreased somatosensory functions, such as mechano- and warmth sensation. Molecular analysis revealed the effect of ePLs for increasing PIEZO mechanosensitivity and lowering the temperature threshold for TRPA1 activation. Finally, we analyzed the effects of ePLs on the physicochemical properties of cell membranes and found that ePLs altered membrane tension and lipid order. Our study demonstrates that ePLs play pivotal roles in optimizing multiple somatosensory modalities by altering cell membrane properties and tuning functions of distinct receptors.

## RESULTS

### Ether phospholipids are enriched in Drosophila neurons

Given the limited number of studies reporting ePLs synthesis in *Drosophila melanogaster*, we first confirmed the presence of ePLs in different tissues. We dissected various tissues from third instar larvae or used the whole body to quantify mRNA expression level of *AGPS* (Figure 1A). *AGPS* was abundantly expressed in the CNS (4.5 times more than in whole body), whereas trace levels of expression were observed in the fat body and carcass (Figure 1A). *AGPS* expression was also analyzed in peripheral sensory neurons in the body wall. Using magnetic bead-based cell sorting, we isolated *ppk*-positive sensory neurons, a subset of polymodal, class IV multidendrtic neurons^27^. The *AGPS* expression level in the magnetic bead-bound (class IV neuron enriched) fraction was significantly higher than in the bead unbound (other cells from the body wall) fraction (p < 0.01 using Student’s t-test, Figure 1B). Consistent with the ePL abundance in neuronal tissues, third instar larvae carrying *AGPS CRIMIC-GAL4* and *UAS-GFP* showed strong *AGPS* expression in neuronal tissues, including the CNS and peripheral sensory neurons such as class IV multidendritic neurons (Figures S1A–D). These findings suggest that ePL plays functional roles in central and peripheral neurons in *Drosophila*.

**Figure 1.**
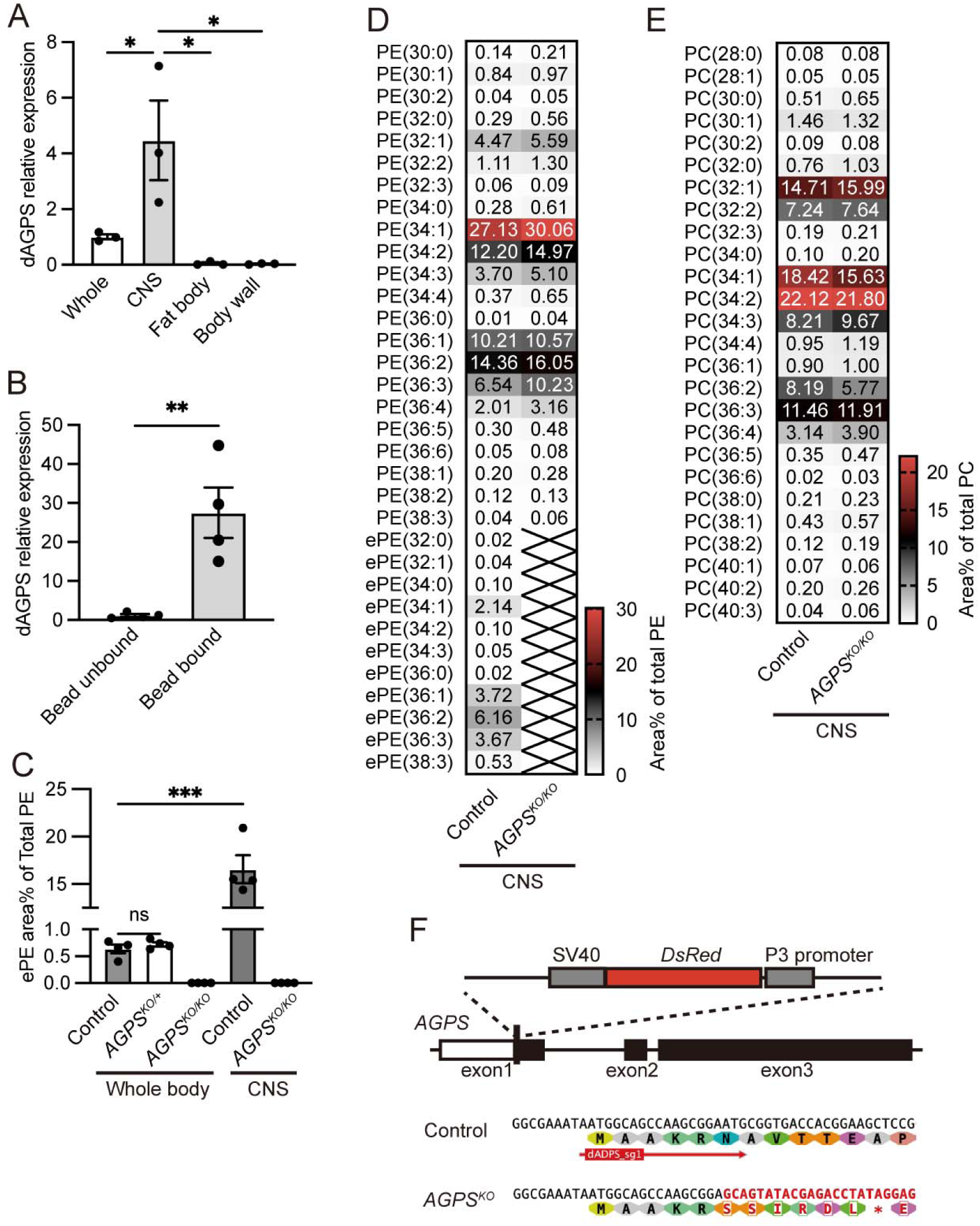
Ether phospholipids are enriched in *Drosophila* neurons. **A** Relative *AGPS* expression level in different tissues in *w^1118^* third instar larvae (n = 3). CNS: central nervous system. *p < 0.05; Tukey’s test. **B** Relative *AGPS* expression level in *ppk-*positive peripheral neurons (bead-bound) and other cells (bead unbound) (n = 4). ** p < 0.01; Student’s t-test. **C** Total ePE area (% of total PE in whole body and the CNS) of third instar larvae (n = 4) in *w^1118^* (control), heterozygous *AGPS* knockout (*AGPS^KO/+^*), and homozygous *AGPS* knockout (*AGPS^KO/KO^*) are shown. *** p < 0.001; Student’s t-test. Each point in **A**–**C** represents a biological replicate. Data are presented as mean ± SEM. **D, E.** Colormap for the composition of PE (**D**) and PC (**E**) species (area% of total PE or PC) in the CNS of *w^1118^* (control) and homozygous *AGPS^KO^* third instar larvae (n = 4). Phospholipid molecules are listed as PE (X:Y) or PC (X:Y), where X and Y denote the total number of carbon molecules in acyl chains and double bonds in acyl chains, respectively. Cross marks indicate phospholipid species under detectable levels. **F** Genomic structure of *Drosophila AGPS^KO^*. The *DsRed* gene was inversely inserted into a CRISPR-Cas9 cleaved site downstream of the start codon in the first exon of *AGPS*. The aberrant stop codon in *AGPS^KO^*is indicated by an asterisk.

Next, to determine ePL composition in *Drosophila* membrane phospholipids, we performed a comprehensive lipid analysis of phospholipid species, including ePLs, in the whole body and the CNS. Specifically, two major classes of phospholipids in *Drosophila*^28^, PE and phosphatidylcholine (PC), were analyzed (Figures 1C–E, S1E, F, Tables S1–4). The proportion of ePEs in total PEs was 0.64% ± 0.08% in the whole body and 16.6% ± 1.5% in the CNS (Figure 1C, D). In contrast, only trace level of ether PCs (ePCs) were observed in the PC for the whole body and the CNS (Figure 1E and Tables S1–4). To confirm the functional roles of AGPS in ePL synthesis, we established *AGPS* knockouts (*AGPS* KO) using CRISPR-Cas9 (Figure 1F). Homozygous *AGPS* KO flies (*AGPS^KO^*) expressed a negligible level of *AGPS* mRNA (Figure S1G), not showing obvious growth defects, such as a delay in pupariation timing (Figure S1H). The lipid analysis clearly demonstrated complete depletion of ePL molecules in the whole body and the CNS of *AGPS^KO^* (Figures 1C–E, S1E, F, Tables S1–4), emphasizing the essential role of *AGPS* in ePL synthesis in *Drosophila*. *AGPS* heterozygous KO flies (*AGPS^KO/+^*) showed a significant reduction in *AGPS* expression level (p < 0.001 using Student’s t-test, Figure S1C) but no changes in the total amount of ePLs compared with control (Figures 1C, S1E, F, Tables S1, 2). It has been reported that *AGPS* is expressed both in glial and neural cells in mammals^29^. To determine the cellular source of ePL biosynthesis in the CNS, we performed tissue-specific *AGPS* knockdown using pan-neuronal (*elav-GAL4*) and pan-glial (*repo-GAL4*) drivers. Pan-neuronal *AGPS* knockdown significantly reduced ePE levels in the CNS, while pan-glial knockdown had no effect (Figures S1 I–K, Tables S5, 6), indicating that AGPS-mediated ePL biosynthesis occurs predominantly in neurons.

In mammals, the major ePL molecule is the plasmalogen type, where an *sn-*1 acyl chain is attached to the glycerol backbone via a vinyl–ether bond. To identify the *sn*-1 and *sn*-2 acyl chain profiles of ePEs in *Drosophila*, we performed product ion scan analysis for major ePEs, including ePE (34:1), ePE (36:1), ePE (36:2), and ePE (36:3), in the CNS sample (Figures S2). ePE (36:2) appeared to be the most abundant among all ePEs in *Drosophila*’s CNS (Figure 1D). The major product ion of ePE (36:2) coincided with linoleic acid (C18:2) (Figure S2H), indicating that the *sn*-2 ester-bound acyl group is C18:2, and the ether-linked *sn*-1 hydrocarbon chain was octadecyl alcohol without double bonds (O-18:0). The other analyzed ePEs harbored either palmitoleic acid (C16:1), oleic acid (C18:1), or linolenic acid (C18:3) at the ester-linked acyl chain and O-18:0 at the ether-linked acyl chain (Figures S2B, E, J).

In conclusion, ePLs, especially ePEs, are abundantly present in *Drosophila* neurons. Moreover, ePEs in *Drosophila* neurons appear to contain an ether bond, but not a vinyl–ether bond, at the *sn-1* position, in contrast to the most common ePL molecule found in mammals.

### AGPS mutation alters somatosensory responses in Drosophila

To address the functional roles of ePLs on somatosensation, we analyzed mechano- and thermo-sensory responses in larvae. A recent study demonstrate that linoleic acid supplementation improves PIEZO channel dysfunction and mechanical deficits in Angelman Syndrome^10^. Therefore, we evaluated mechanosensory responses in larvae with *AGPS* mutation and associated ePL loss. First, we assessed tactile sensation by measuring responses to a gentle touch to the nose of a freely moving late third instar larva (120 h after egg laying, AEL); no difference in responses was observed between control and *AGPS^KO^* larvae (Figure 2A). Second, we characterized mechanical nociception by poking a larva using von Frey filaments of three different forces (20.1, 47.0, and 70.0 mN). Nociceptive rolling behavior to the weakest force (20.1 mN) was significantly decreased, and decreasing tendency to stronger ones (47.0 mN) was observed in *AGPS^KO^* compared with control larvae (p < 0.05 using Student’s t-test, Figure 2B). Reduced response to von Frey filament stimulation reminiscent of defects observed in PIEZO knockout larvae^30^, although it was substantially milder. These results demonstrated that ePL loss leads to an abnormal mechanical nociceptive phenotype in *Drosophila*.

**Figure 2.**
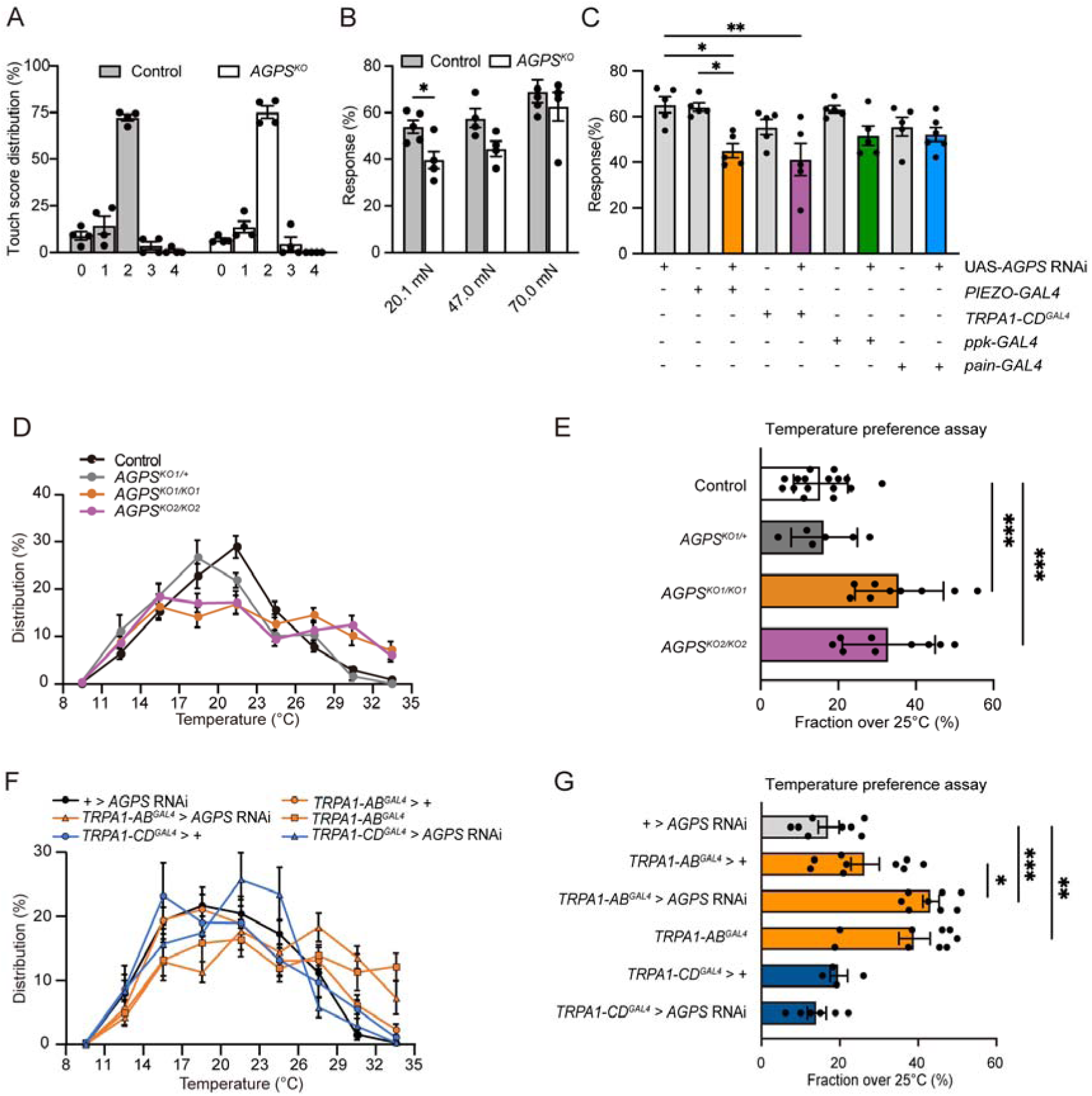
AGPS influences multiple sensory modalities in *Drosophila*. **A** Distribution of scored responses to gentle touch with a nickel–titanium filament in *w^1118^* (control) and *AGPS^KO^*late third instar larvae (n = 4). 0: no response, 1: hesitation, 2: turn or withdrawal of the anterior segments, 3: single reverse contractile, or 4: multiple reverse contraction. **B** Responses to different forces applied with von Frey filaments (20.1, 47.0, and 70.0 mN) in *w^1118^* (control, n = 4 or 5) and *AGPS^KO^* late third instar larvae (n = 4 or 5). **C** Responses to a von Frey filament (20.1 mN) in UAS control (*UAS-AGPS RNAi*/+, n = 5), GAL4 controls (*PIEZO-GAL4;* n = 5, *TRPA1-CD^GAL4^*; n = 5, *ppk-GAL4;* n = 5, and *pain-GAL4*; n = 5), and *AGPS* knockdown larvae using *PIEZO-GAL4* (n = 5), *TRPA1-CD^GAL4^* (n = 5), *ppk-GAL4* (n = 5), and *pain-GAL4* (n = 6). Data are presented as mean ± SEM. *p < 0.05, **p < 0.01; Student’s t-test (**B**) or Tukey’s test (**C**). **D**, **E** Temperature preference assay on a linear gradient for late third instar larvae of *w^1118^* (control, n = 16), heterozygous *AGPS^KO^* (*AGPS^KO1/+^*, n = 6), and homozygous *AGPS^KO^* clones (*AGPS^KO1/KO1^* and *AGPS^KO2/KO2^*, n = 9). **F**, **G** Temperature preference assay for late third instar larvae of UAS control (*UAS-AGPS RNAi*/+, n = 8), GAL4 controls (*UAS-dicer-2*/+;*TRPA1-AB^GAL4^*/+, n = 9 and *UAS-dicer-2*/+;*TRPA1-CD^GAL4^*/+, n = 4), *TRPA1-AB* knockout (*TRPA1-AB^GAL4^*/*TRPA1-AB^GAL4^*, n = 9), *AGPS* knockdown by *TRPA1-AB^GAL4^* (*UAS-dicer-2/UAS-AGPS* RNAi;*TRPA1-AB^GAL4^*/+, n = 8) and *TRPA1-CD^GAL4^* (*dicer-2/AGPS* RNAi;*TRPA1-CD^GAL4^*/+, n = 6). Distribution of third instar larvae on a thermal gradient of 8°C–35°C (**D, F**) and the fraction of larvae distributed over 26°C zones (**E, G**). Data are presented as mean ± SEM. *p < 0.05; **p < 0.01, *** p < 0.001; Dunnett’s test or Tukey’s test.

To investigate whether sensory functions are affected by the absence of ePLs, we used the GAL4-UAS system for tissue specific *AGPS* knockdown in mechanosensory neurons that express PIEZO channels involved in mechanical nociception. Several receptor genes are considered responsible for von Frey filament responses, including *PIEZO*^30^, *TRPA1-CD*^31,32^, *ppk*^33^, and *painless*^34^. When *AGPS* RNAi was induced using *PIEZO-GAL4*, we observed a significant decrease in the response to the von Frey filament compared to both UAS and GAL4 controls (20.1 mN, p < 0.05 using Tukey’s test; Figure 2C) and *TRPA1-CD^GAL4^* induced RNAi conferred significant decrease in the response of von Frey filament compared to UAS control (20.1 mN, p < 0.01 using Tukey’s test; Figure 2C). In *PIEZO-GAL4*-dependent *AGPS* knockdown, we observed a significant decrease in the response to the von Frey filament compared to both RNAi and *PIEZO-GAL4* controls (20.1 mN, p < 0.05 using Tukey’s test; Figure 2C). *TRPA1-CD^GAL4^*-induced *AGPS* knockdown was not significantly lower than *TRPA1-CD^GAL4/+^*due to its heterozygote phenotype, but the trend toward additional impairment supports our conclusions. These results suggest that AGPS and its product, ePLs, play a crucial role in modulating the mechanosensory responses of PIEZO-expressing neurons in *Drosophila*.

A previous study reported that alterations in membrane lipid composition can affect temperature preference in *Drosophila*^19^. Therefore, we next investigated the effect of ePLs on thermosensation in larvae. We subjected late third instar larvae to a temperature gradient assay (8°C–35°C) to examine their thermotactic behavior (Figures S3A, B). Control (*w^1118^*) larvae distributed through the temperature gradient with highest proportions in 20°C–23°C zone, avoiding lower and higher temperatures (Figure 2D). However, *AGPS* KO (*AGPS^KO^*) lost a clear peak in their distribution, showing increased fractions in high temperature zones than the control larvae (Figure 2D). Specifically, the proportion of larvae distributed over 26°C zones was 15.5% ± 1.8% in control, whereas significant increases (> 30%) were observed in *AGPS^KO^* larvae (35.8% ± 3.8%; p < 0.001 using the Dunnett’s test, Figure 2E).

We then used different GAL4 lines for *AGPS* knockdown to identify the thermosensory neurons involved in the altered thermotaxis. We initially used GAL4 lines labeling TRPA1-positive neurons involved in warm temperature avoidance^35,36^. In *Drosophila*, there are four functional TRPA1 splicing variants transcribed from two distinct promoters (TRPA1-AB and TRPA1-CD)^31,37^. Among them, *TRPA1-AB^GAL4^*-induced *AGPS* knockdown resulted in a significant increase in the fraction of larvae distributed in high temperature zones compared to the controls (p < 0.001 vs. *AGPS* RNAi control, p < 0.05 vs. *TRPA1-AB^GAL4^* control, using Tukey’s test, Figures 2F, G), which resembled the *AGPS^KO^* phenotype (Figures 2D, E). *TRPA1-AB^GAL4/+^* showed a slightly higher fraction over 26°C because this line represents a heterozygous knockout of *TRPA1-AB* (GAL4 insertion disrupts the AB isoform), but *AGPS* knockdown in these neurons exhibited significantly higher fractions compared to the GAL4 heterozygotes and the RNAi control, indicating a genuine genetic interaction rather than simple additivity (p > 0.05, using Tukey’s test Figures 2F, G). We also tested GAL4 lines labeling TRPL-^38^ and Inactive (Iav)-positive neurons^39^, involved in cool temperature avoidance. However, no significant changes in either warm or cold temperature avoidance were observed (Figures S3C, D, E).

To further confirm that TRPA1-dependent thermotactic behaviors depend on *AGPS*, larvae were subjected to a thermal two-choice assay (20°C vs. 29°C). The higher temperature was highly aversive for control larvae, whereas reduced or no avoidance was observed in *AGPS^KO^*(Figures S3F) or *AGPS* knockdown in TRPA1-AB neurons (Figure S3G), respectively. The avoidance index of *TRPA1-AB^GAL4/+^* control was significantly lower than both wild-type and *UAS-AGPS RNAi/+* controls (Figure S3G), reflecting its nature as a *TRPA1-AB* heterozygous knockout. Notably, TRPA1-AB-specific *AGPS* knockdown produced more severe thermal avoidance defects than whole-animal *AGPS* knockout, likely because tissue-specific disruption avoids potential compensatory interactions between multiple altered sensory systems that occur in global knockouts. Taken together, these results suggest that AGPS and its product, ePLs, could modulate responsiveness of both thermo- and mechanosensory neurons.

To further confirm the involvement of ePLs in somatosensory responses, we tested whether behavioral abnormalities in *AGPS^KO^* larvae could be restored by the dietary supplementation of octadecanoyl glycerol (18-AG), an ePL precursor. The results revealed differential rescue effects between sensory modalities. In mechanical nociception assay, 18-AG supplementation showed a trend toward restoration of nociceptive responses to von Frey filament (20.1 mN) in *AGPS^KO^* larvae, although this did not reach statistical significance (p=0.13 vs. *AGPS^KO^* using Tukey’s test) (Figure S3H). On the other hand, in thermal gradient assay, 18-AG supplementation failed to ameliorate the abnormal *AGPS^KO^* larvae in high-temperature zones (Figures S3I, J). This differential pattern is consistent with previous reports showing that dietary alkylglycerol supplementation increases ePL levels in peripheral tissues but not in the CNS^40^. These findings suggest that endogenous ePL biosynthesis is particularly crucial in the CNS for maintaining normal sensory function in *Drosophila*.

As it has been reported that phospholipid homeostasis affects the morphogenesis of sensory neurons^41,42^, we assessed the morphology of sensory neurons. Peripheral multidendritic neurons expressing PIEZO channels showed no obvious morphological changes following *AGPS* knock down (Figures S3K, L). Similarly, the morphology of warm sensitive TRPA1-A/B-expressing neurons in parts of the CNS in larvae^35^ was not affected by *AGPS* knockdown (Figures S3M–P), although quantitative analysis was precluded by the deep location of these neurons and consequent variability in immunostaining penetration. These results suggested that *AGPS* reduction may not alter the development and cell fate of these sensory neurons.

### Responsiveness of PIEZO is enhanced by ether phospholipids

As membrane lipid composition affects the ion channel properties, including PIEZO and TRP channels, we hypothesized that AGPS-produced ePLs influence somatosensation in larvae by directly modulating the activity of the somatosensory channel proteins, such as PIEZO and TRPA1. To elucidate the functional role of ePLs in channel proteins, we conducted an electrophysiological analysis using cultured cells. We utilized S2R+ cells, a *Drosophila* embryonic cultured cell line that lacks the ability to synthesize ePLs. To establish ePL-producing S2R+ cells, we employed supplementation of alkylglycerol (AG) as a precursor of ePLs to wild type cells to bypass the requirement of AGPS to drive ePL biosynthesis (Figure 3A).

**Figure 3.**
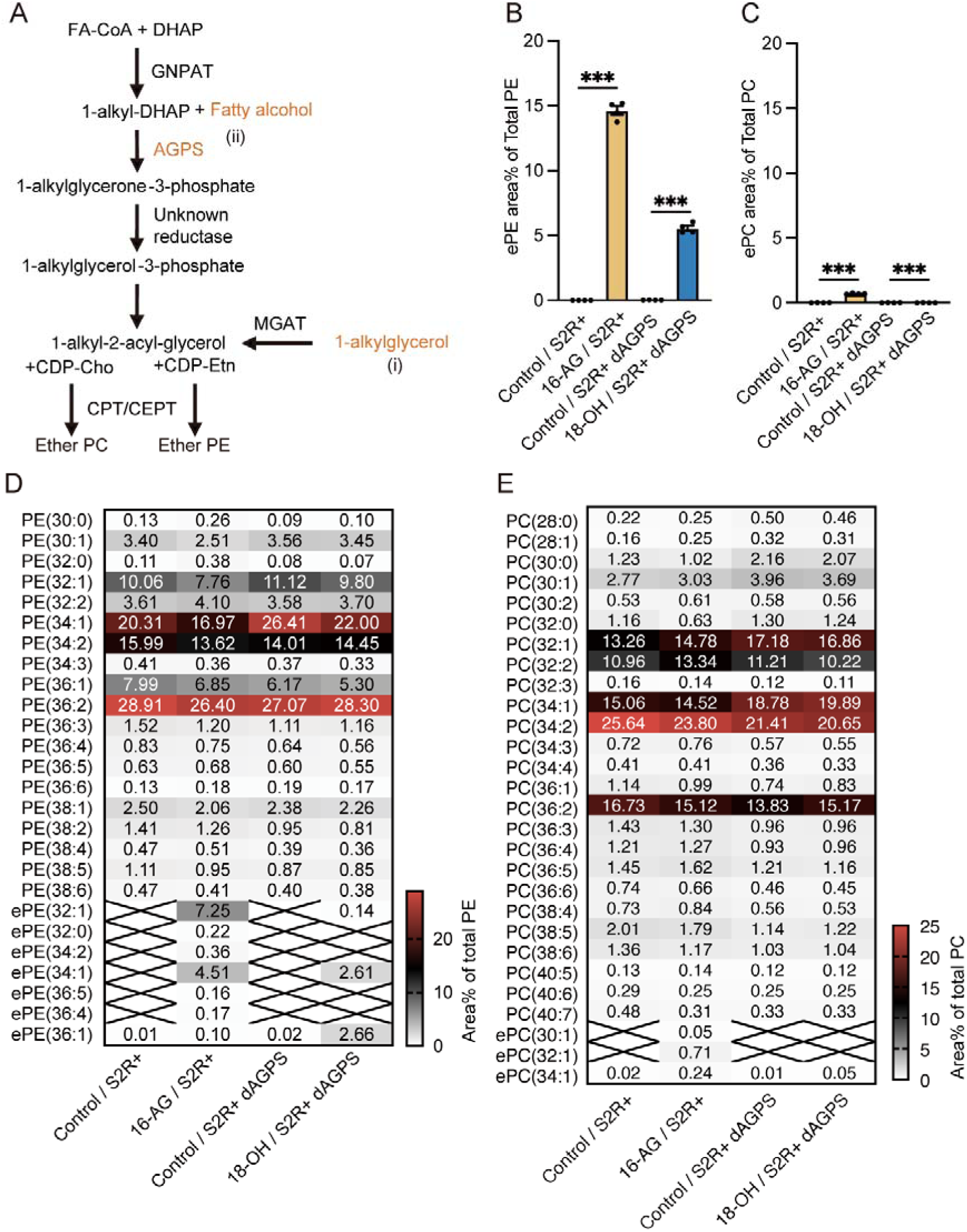
Establishment of ePL-producing S2R+ cells. **A** Synthesis pathway of ePLs. Two strategies to increase ePL production in *Drosophila* are shown. (i) Supplementation with 16-AG or 18-AG (1-alkylglycerol) or (ii) *AGPS* overexpression with 18-OH (fatty alcohol) supplementation. FA-CoA, Fatty acyl-CoA; DHAP, dihydroxyacetone phosphate; GNPAT, glyceronephosphate O-acyltransferase; MGAT, monoacylglycerol acyltransferase; CDP, cytidine diphosphate; CPT, choline phosphotransferase; and CEPT, choline/ethanolamine phosphotransferase. **B–E** Lipid composition in control and ePL-producing S2R+ cells (n = 4) for total ePE area% of total PE (**B**), total ePC area% of total PC (**C**), and colormap for composition of PE (**D**) and PC (**E**) species (area% of total PE or PC) in control cells (Control/S2R+), 100 µM 16-AG supplemented cells (16-AG/S2R+), dAGPS-expressing cells (Control/S2R+ dAGPS), and 100 µM 18-OH supplemented *dAGPS*-expressing cells (18-OH/S2R+ dAGPS). Phospholipid molecules are listed as PE (X:Y) or PC (X:Y), where X and Y denote the total number of acyl chains and double bonds in acyl chains, respectively. Cross marks in (**D, E**) indicate phospholipid species under detectable levels. Each point represents a biological replicate. Data are presented as mean ± SEM. ***p < 0.01; Student’s t-test.

The lipid analysis revealed successful ePL production in S2R+ cells with alkylglycerol supplementation (Figures 3B–E, Tables S7, 8). Importantly, the production of ePEs was primarily induced, consistent with the ePL composition in larvae (Figures 1D, E). A product ion scan analysis demonstrated that supplementation with alkyl glycerol (16-AG) induced the production of ePE (32:1) and ePE (34:1), considered to introduce palmitoleic acid (C16:1) and oleic acid (C18:1) at the *sn-*2 ester-linked position of ePE, respectively (Figures S4A, B), suggesting that a acyl alcohol without double bonds was bound at the ether-linked position. Thus, the composition of the *sn-*1 hydrocarbon chain in ePLs appears to reflect the features of the supplemented AG or fatty alcohol.

Difference in membrane lipid composition reportedly leads to alterations in membrane properties, thereby affecting the activity of mechanosensory PIEZO channels^9,10^. To test the functional interaction between ePLs and PIEZO, we first evaluated PIEZO response in S2R+ cells to its activator Yoda1^43^ in the absence or presence of 18-AG, which is the major *sn-*1 hydrocarbon chain in ePLs (Figure S2) in a Ca^2+^ imaging. Ca^2+^ ionophore ionomycin was applied after Yoda1 to calibrate the maximum capacity of Ca^2+^ intake. We set a stringent threshold (Δratio > 2) to exclude non-specific calcium fluctuations from sources other than PIEZO activation. PIEZO-expressing S2R+ cells were stimulated with continuous perfusion of Yoda1, which increased the proportion of cells showing robust calcium responses, suggesting enhanced PIEZO sensitivity to Yoda1 stimulation (11.1% ± 3.1%; Figures 4A, B, D, E). When S2R+ cells were supplemented with 18-AG, Yoda1 perfusion significantly elicited higher Ca^2+^ increases in a larger cell population (28.7% ± 3.8%, p < 0.01 using Student’s t-test; Figures 4A, C–E). Next, we performed electrophysiological recordings of PIEZO activation using mechanical stimuli in PIEZO-overexpressing S2R+ cells. Due to the small size and non-adherent properties of S2R+ cells, we used a single mechanical stimulus (6-μm deflection depth) to assess PIEZO mechanosensitivity and avoid potential cell damage from deeper deflections. A poking-induced small activation, followed by rapid desensitization was observed in PIEZO-overexpressing cells (Figures 4F, H). When cells were supplemented with 18-AG, the current density of PIEZO-expressing S2R+ cells significantly increased (p < 0.01 using Mann–Whitney U test, Figures 4G, H). However, the proportion of poking responding cells did not change in the presence/absence of ePLs when we use a sensitive threshold (> 5 pA; Figure 4I). These results suggested that ePLs enhance PIEZO sensitivity, thereby influencing animals’ mechanical responses. These results suggest that ePLs, particularly ePEs in the plasma membrane, enhance the sensitivity of PIEZO channel activation, which may influence mechanical nociception in animals.

**Figure 4.**
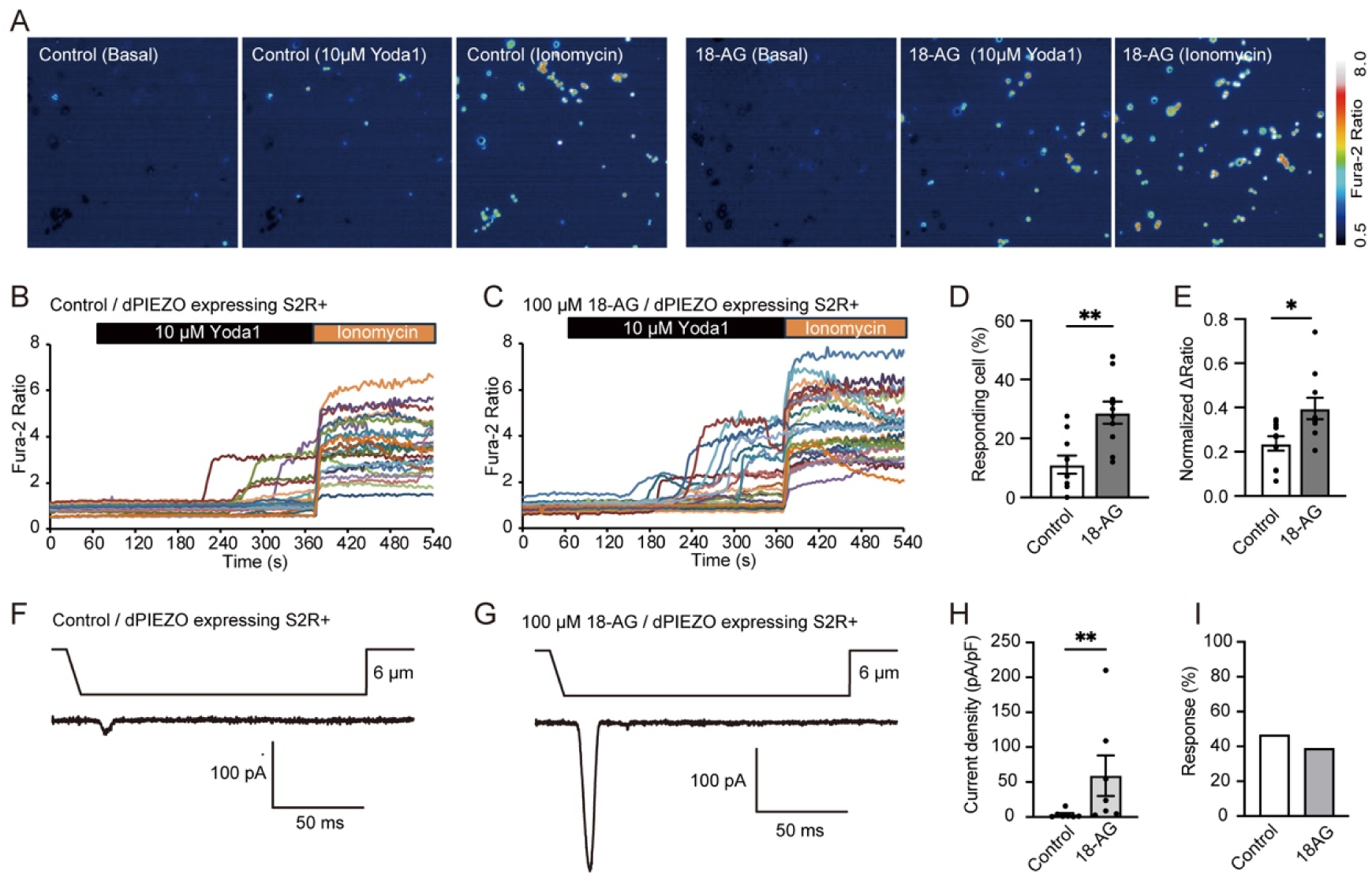
ePLs modify dPIEZO activity in *Drosophila* cells. **A–C** Representative Fura-2 imaging data for PIEZO in control S2R+ cells and cells supplemented with 100 µM 18-AG. **A,** Typical ratiometric images before (Basal, at 60 s) and after application of 10 µM Yoda1 (Yoda1, at 360 s) or 5 μM ionomycin (Ionomycin, at 540 s) in dPIEZO expressing cells (left) and dPIEZO expressing cells supplemented with 100 μM 18-AG (right). **B, C.** Representative Ca^2+^ level traces in dPIEZO expressing cells (**B**) and 18-AG supplemented dPIEZO expressing cells (**C**). **D, E** Proportion of cells responding to Yoda1 (Δratio > 2) (**D**) and maximum Δratio response to Yoda1 normalized by the ionomycin response (**E**) in dPIEZO expressing cells (control, n = 10) and 18-AG supplemented dPIEZO expressing cells (18-AG, n = 10). Each point represents a biological replicate; 25―40 cells were analyzed in each assay. Data are presented as mean ± SEM. *p < 0.05, **p < 0.01; Student’s t-test. **F, G** Representative traces of patch-clamp recordings of mechanical stimuli-evoked dPIEZO activation without (**F**) or with (**G**) 100 μM 18-AG. **H** Quantification of the peak current density. Data are presented as mean ± SEM. **p < 0.01; Mann–Whitney U test. **I** Proportion of responding cells (current size > 5 pA).

### Ether phospholipids potentially reduce the temperature threshold for dTRPA1 activation

Next, we evaluated the temperature-dependent dTRPA1-A activation using S2R+ cells. The temperature threshold for dTRPA1-A activation was 22.1°C ± 0.4°C in control S2R+ cells (Figures 5A, C), significantly decreasing (p < 0.01 using Student’s t-test) to 20.8°C ± 0.3°C when S2R+ cells were supplemented with 18-AG (Figures 5B, C). In contrast, we did not observe alterations in maximum current densities due to heat or electrophilic TRPA1 activator, allyl isothiocyanate (AITC), stimulation in ePL-producing cells (Figures 5D–F). We also assessed the temperature-dependent dTRPA1-A activation in another ePL producing cells overexpressing AGPS with fatty alcohol supplementation (Figure 3A). Supplementation with octadecyl alcohol (18-OH) to AGPS-expressing cells resulted in the production of ePE octadecyl alcohol (O-18:0) at the ether-linked positions (Figure S4C). Consistent with the 18-AG supplementation, 18-OH supplementation to AGPS-expressing S2R+ cells significantly decreased the temperature threshold for dTRPA1-A activation compared with that for control cells (20.3°C ± 0.7°C vs. 24.0°C ± 1.2°C; p < 0.05 using Student’s t-test, Figures S5A–C). Likewise, neither heat-nor AITC-evoked current densities were affected (Figures S5D–F). Because the *TRPA1-AB^GAL4^*-induced *AGPS* knockdown flies showed a significant decrease in warmth avoidance (Figures 2F, G), dTRPA1-B properties were also assessed in ePL-producing S2R+ cells, whose temperature sensitivity has been reported previously^37^. Both the temperature threshold for activation and the maximum heat-evoked current densities were unaltered in the presence of ePLs (Figures S5G–J). In contrast, AITC-induced current density was significantly reduced in ePL-producing S2R+ cells at low (30 µM) and high (300 µM) concentrations (p < 0.05 using Student’s t-test, Figures S5K, L). In addition, the deficient in thermosensitive rolling behavior of *TRPA1-KO* flies restored by rescue expression of dTRPA1-A but not of dTRPA1-B isoform ^35^. Thus, dTRPA1-B and its expressing neuron may not involve in the ePL-dependent thermoregulatory behavior. We also confirmed no thermal activation was observed in the non-transfected S2R+ cells (Figure S5M), confirming that the thermal responses observed were specifically derived from transfected TRPA1 isoforms rather than endogenous cellular mechanisms (Figures 5A, B, S5A, B, G, H).

**Figure 5.**
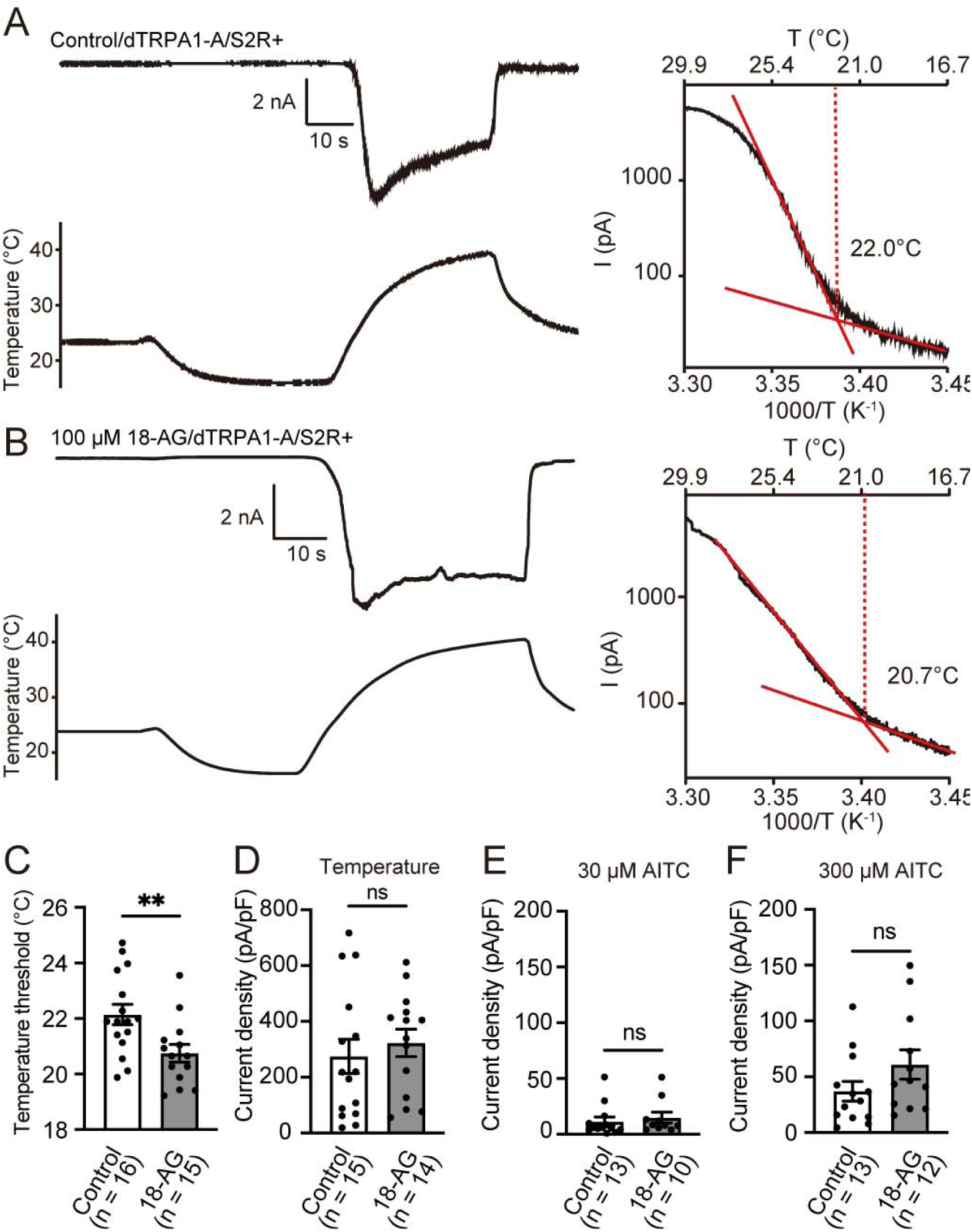
ePLs modulate the temperature threshold for dTRPA1-A activation. **A, B** Whole-cell patch-clamp recording of dTRPA1 activation with temperature stimulation in dTRPA1 expressing S2R+ cells (**A**), dTRPA1 expressing cells supplemented with 100 µM 18-AG (**B**), Left, representative traces of recordings; right, Arrhenius plots from the traces in the left panels. Temperature thresholds were determined as a crossing point of two fitted lines. **C–F** Quantification of the temperature threshold (**C**) and peak current densities from heat (**D**), 30 µM AITC (**E**), and 300 µM AITC (**F**) stimulation in dTRPA1 expressing cells supplemented with solvent (control) and 18-AG. Each point represents a biological replicate. The number of replicates (n) is shown in the panels. Data are presented as mean ± SEM. **p < 0.01; Student’s t-test.

Taken together, these results suggest that ePLs, particularly ePEs in the plasma membrane, reduce the temperature threshold for heat-evoked dTRPA1-A activation, which may partially explain the thermotaxis defects in *AGPS^KO^* larvae.

### Ether phospholipids alter membrane tension and lipid order

Previous reports have shown that lipid modulation alters membrane tension, which can impact the activity of sensory channels^9,44^. To get further insights into the effects of ePEs on membrane properties in S2R+ cells, we used different approaches to quantify membrane tension and lipid order.

First, we measured membrane tension through atomic force microscopy (AFM) force curve measurement. Parts of the cell surfaces were scanned with AFM while intermittently recording force curves to determine the Young’s modulus, which reflects membrane tension at each point (Figure S6A). The median value of the calculated Young’s modulus in control was 0.95 × 10^5^ Pa (Figure 6A) while that in 18-AG supplemented S2R+ cells was 2.4 × 10^5^ Pa, significantly higher (p < 0.001, Mann–Whitney U test) than control cells. Second, we utilized a Flipper-TR probe to measure membrane tension. When the probe is incorporated into the plasma membrane, the emitted fluorescent lifetime correlates with the membrane tension^45^. A significant longer fluorescent lifetime (p < 0.05, Mann–Whitney U test) was observed for 18-AG supplemented S2R+ cells compared with the control (Figures 6B, S6B). These distinct measurements consistently showed an increase in membrane tension in the presence of ePLs in the plasma membrane.

**Figure 6.**
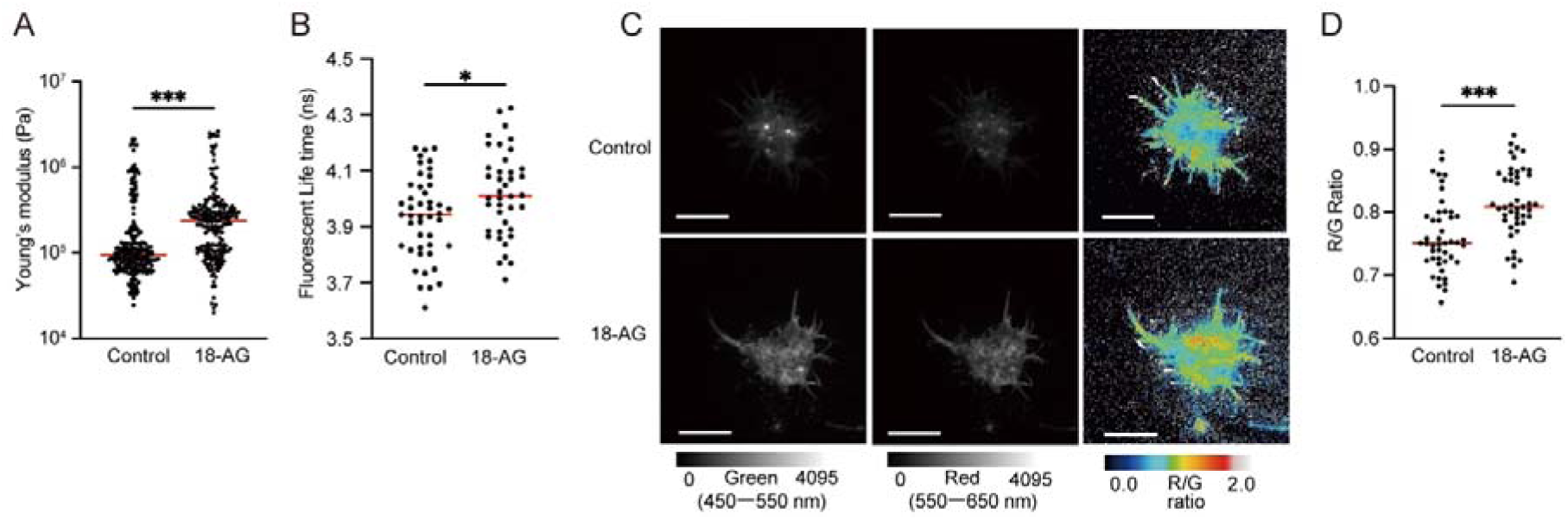
ePLs alter membrane properties of *Drosophila* cells. **A** Measurement of membrane tension using AFM in S2R+ cells (control) and cells supplemented with 100 μM 18-AG. Data were acquired from 242 different regions of 24 control cells and 213 different regions of 27 cells supplemented with 18-AG. The red horizontal lines represent the median and each point is the averaged Young’s modulus (Pa) of each region. **B** Measurement of membrane tension with fluorescence lifetime imaging using Flipper-TR in control (n = 45) and 100 μM 18-AG supplemented S2R+ cells (n = 40). Red horizontal lines represent the median and each point indicates the average fluorescent lifetime of a single cell. **C** Representative pseudocolor images of LipiORDER loaded control (upper) and 100 μM 18-AG supplemented S2R+ cells (lower). Green fluorescence intensity (left, 450–550 nm), red fluorescence intensity (middle, 550–650 nm), and ratiometric images of red to green fluorescence intensity (right, R/G ratio). Scale bars, 100 µm. **D** Quantification of red/green fluorescence ratio (R/G ratio) in control (n = 46) and 100 μM 18-AG supplemented S2R+ cells (n = 45). Each value indicates the average R/G ratio of a single cell. Red horizontal lines represent the median and each point is a biological replicate. Data are presented as mean ± SEM. *p < 0.05, ***p < 0.001; Mann–Whitney U test.

Last, we assessed the lipid order using a LipiORDER Dye^46^. A significant red shift (p < 0.001, Mann–Whitney U test) in fluorescence spectrum was observed in 18-AG supplemented S2R+ cells (Figures 6C–D, S6C), suggesting that ePE-harboring membranes are in more lipid-disordered states than ePE-depleted membranes. We also utilized another lipid order sensitive dye, laurdan^47^ to confirm the effect of 18-AG supplementation on membrane fluidity. Consistent with our LipiORDER results, 18-AG treatment significantly decreased the GP score (p < 0.05, Mann–Whitney U test), confirming increased membrane fluidity in ePL-containing S2R+ cells (Figure S6D)^47^. Taken together, these results demonstrated that ePEs modify the physicochemical properties of the plasma membrane, which potentially leads to changes in the activity and/or sensitivity of sensory proteins, such as PIEZO and TRPA1.

## DISCUSSION

In this study, we identified ePLs as a novel class of lipid species that modulates ion channel activity and somatosensory functions in *Drosophila.* ePLs have received increased attention due to their functional association with neurodegenerative diseases and metabolic disorders accompanied by systemic inflammation; potential roles, such as an oxidative stress scavenger, have been proposed. Moreover, recent studies have explored their involvement in ferroptosis and cancer treatment^48^. Although ePLs are closely linked to several pathological events, physiological targets of ePLs remain unclear. Our data demonstrate that ePLs are required for intrinsic sensory responses by modulating the responses of multiple sensory proteins, such as PIEZO and TRPA1, in the PNS and CNS. The mechanisms of ion channel activity modulation by ePLs are open for discussion. It has been discussed that PUFAs in the *sn-*2 position of ePLs and their metabolites as direct activators of ion channels^49^. Moreover, the ion transport activity of sarcosomal sodium–calcium exchanger in plasmalogen-containing proteoliposome has been reported^25^. Recently, a possible correlation between changes in cell membrane tension due to oxidative stress and the activation of mechanoreceptor channels and cell death has been proposed^50^. These evidences underscore ion channels as a molecular target in the regulation of biological processes by ePLs.

Our study clearly identified the functional roles of nonplasmenyl-type ePLs on sensory channels. Product ion scan analysis of major ePL species in the CNS showed a prevalence of plasmanyl-type ePLs, which lack double bonds, rather than plasmenyl-types harboring a vinyl–ether linkage with a double bond (Figure S2). In humans, approximately 20% of ePLs are plasmenyl-type plasmalogen^21^. Lipidomic analysis of honeybee (*Apis mellifera*) showed abundant nonplasmenyl-type ePLs in the CNS^51^, suggesting the prevalence of plasmanyl-type ePLs in the CNS of insects. To date, most research has focused on the function of plasmenyl-type plasmalogen ePLs, particularly on the high sensitivity of the vinyl–ether linkage with a double bond to reactive oxygen species and its correlation with oxidative stress^52^. Recently, a Δ1-desaturase, plasmanylethanolamine desaturase 1 (PEDS1/TMEM189), has been identified as the enzyme responsible for inserting a double bond at the *sn*-1 position of ePLs^53,54^. However, the putative PEDS1 gene, *Kua*, does not show high expression in the CNS according to FlyAtlas RNA-seq analysis^55^. This could explain the minimal presence of plasmenyl-type ePLs in the CNS of *Drosophila* (Figure S2). It would be interesting to specifically explore the expression pattern and function of *Kua* in *Drosophila*.

The alteration of membrane properties may be one of the key factors for the molecular basis of functional correlation between ePLs and ion channels. Previous research on mouse PIEZO1 showed that an increase in saturated FA in the cell membrane negatively alters channel activity by bringing the membrane to an “ordered” state^9^. Since our LipiORDER analysis revealed that the membrane in ePL-lacking control S2R+ cells exhibited a more “ordered” state than ePL-producing cells (Figure 6C), this may explain the lower activity of *Drosophila* PIEZO in control cells. In addition, our AFM and Flipper-TR imaging-based assays demonstrated that ePLs increase cell membrane tension (Figures 6A, B). As PIEZO responds to changes in cell membrane tension and deformation^56^, the increased activity observed in ePL-producing S2R+ cells may be due to altered force thresholds for inducing cell membrane deformation. Regarding PIEZO mechanosensitivity, our AFM and Flipper-TR analyses demonstrate that ePLs increase membrane tension (Figures 6A, B). While this appears contradictory to previous studies showing that membrane stiffening (via saturated fatty acids) reduces PIEZO activity^9^, we propose that ePL-mediated stiffening has distinct mechanistic effects. We hypothesize that ePLs promote lipid microdomain formation, particularly under increased membrane tension. Studies have shown that elevated tension enhances microdomain formation in large model membranes^57–59^, and Flipper-TR may detect stiffness changes associated with such domains^45^. PIEZO activation requires structural transitions from curved (closed) to flattened (open) conformations during membrane extension^56,60^. ePLs may facilitate PIEZO localization to specific lipid microdomains that stabilize the resting curved state, thereby enhancing sensitivity to membrane deformation. Critically, PIEZO sensitization occurs when resting membrane tension is minimized, allowing greater dynamic range for activation^61^. ePL-induced compartmentalization may create local environments that optimize this tension differential^9,56,60,61^.

For TRPA1 thermosensitivity, the relationship between membrane stiffness and temperature sensitivity remains poorly understood for thermosensitive TRP channels. We propose that ePLs may influence temperature-dependent membrane phase transitions and separation^57,59^, creating local lipid environments that lower the energetic barrier for TRPA1 activation. Unlike global membrane stiffening, ePL-mediated changes may create heterogeneous membrane domains with distinct thermodynamic properties. Since ePL’s effect on thermal sensitivity is specific to TRPA1-A but not B isoform (Figures 5, S5), it might rely on the structural difference between A and B isoforms in the linker domain connecting the N-terminus and transmembrane domains^37^. Current macroscopic membrane measurements cannot resolve the spatiotemporal dynamics of local membrane tension and lipid-protein interactions^45^. Future studies using advanced biophysical approaches will be essential to understand how ePLs create functionally distinct lipid environments for different mechanosensory and thermosensory channels. Previous research has shown that eicosapentaenoic acid and its derivatives modulate mammalian thermosensitive TRPV4 responses to ligands through alterations in membrane tension^44^, but whether mechanical and temperature activation of TRPV4 is affected remains unknown. As temperature affects lipid order, differences in lipid order between control and ePL-producing S2R+ cells could be an underlying mechanism for the altered temperature threshold of *Drosophila* TRPA1. For example, a previous report claimed that increasing unsaturated FAs in the cell membrane, which should increase lipid disorder, resulted in enhanced heat responsiveness of TRPA1 neurons and decreased preferred temperature of *Drosophila* larvae^19^. Further research is needed to clarify the cell membrane properties and their underlying mechanisms that modulate the temperature threshold of TRPA1 activation.

It should be noted that our thermosensory measurements of TRPA1 activation, while showing statistically significant effects on thermal threshold reduction, have important limitations that prevent definitive mechanistic conclusions. The variability in absolute thermal thresholds between experimental preparations and the modest effect sizes (∼2°C) relative to behavioral phenotypes indicate that our current in vitro approach may not adequately recapitulate the physiological conditions relevant for thermotaxis. Particularly, the quantitative relationship between ePL-mediated changes in TRPA1 properties and thermotactic behavior requires careful interpretation. Our patch-clamp measurements were performed under rapid heating conditions necessary for reliable threshold determination, show modest but consistent effects of ePL manipulation. However, the thermal threshold change (∼2°C reduction in the presence of ePLs) does not fully explain the thermotaxis defects by AGPS mutation. TRPA1 responses are known to be highly rate-dependent^35^, and we propose that ePL depletion has more pronounced effects on channel responses to the gradual temperature changes characteristic of thermotactic behavior. This rate-dependency could at least explain why small threshold changes measured under rapid heating translate to severe defects in temperature discrimination under natural conditions. Future studies should aim at clarifying how TRPA1-AB neurons react to temperature conditions relevant for thermotactic behavior in the presence or absence of ePLs.

An intriguing aspect of our findings is that ePLs, particularly ePEs, increase membrane stiffness/tension while increasing its fluidity. We propose that this apparent paradox may reflect the complex, heterogeneous nature of ePL-containing membranes.

Several mechanisms could reconcile these observations: First, ePLs may create spatially distinct lipid microdomains with different biophysical properties. The unusual ether linkage and unsaturated fatty acid chains of ePLs could promote the formation of specialized membrane domains that coexist with conventional phospholipid regions. Our macroscopic measurements (whole-cell fluorescence averaging and large AFM tip sampling) would detect the net effects of these heterogeneous domains, potentially masking local variations in membrane properties. Second, ePL-induced changes may differentially affect membrane parameters at different length scales. While local lipid packing may become more disordered (increased fluidity), the overall membrane architecture could simultaneously experience increased tension due to altered lipid-protein interactions or membrane curvature effects. Importantly, our current experimental approaches lack the spatiotemporal resolution to distinguish these potential microdomain effects. Future studies must evaluate the spatial organization and local biophysical properties of ePL-containing membranes.

In conclusion, we showed that ePLs modulate distinct functions of sensory proteins, such as PIEZO and TRPA1, thereby optimizing animals’ sensory responses. These channels are expressed not only in sensory neurons but also in various tissues involved in different physiological functions^62,63^. In addition, ePLs are abundantly present in glial cells in mammals^64^. Since the temperature-dependent motility of microglia is modulated by thermosensitive TRPV4^65^ and PIEZO1 activity is reportedly essential for the myelination^66^, it would be interesting to test whether ePLs are involved in the regulation of these channels in glial cells. Exploring the integrative roles of ePLs as a modulator of receptor proteins may shed light on the various biological processes and diseases related to channel proteins.

## Methods

### Fly stocks and husbandry

*w^111^*^8^ was used as a control unless otherwise specified. The following strains were obtained from Bloomington Drosophila Stock Center (BL) or Vienna Drosophila RNAi Center (VDRC): 5×*UAS-mCD8::GFP* (BL #5137), *piezo-GAL4* (BL #59266), *painless-GAL4* (BL #27894), *AGPS-CRIMIC-Gal4* (BL #79234), *UAS-dicer-2* on II (VDRC #60008), *UAS-dicer-2* on III (VDRC #60009), and *UAS-AGPS* RNAi (VDRC #3321). The following stocks were provided by the indicated investigators: *TRPA1-AB^GAL4^*, *TRPA1-CD^GAL4^*, *iav-GAL4*, and *TRPL-GAL4* (C. Montell) and *ppk-GAL4* (D. N. Cox). All fly strains were outcrossed to the control line (*w^1118^*) for 4–5 generations. Flies were reared on standard cornmeal-yeast medium (19 g agar, 180 g cornmeal, 100 g dry brewer’s yeast Ebios, 250 g glucose, 24 mL Methyl 4-hydroxybenzoate [10% w/v solution in 70% ethanol], and 8 mL propionic acid in 2500 mL reverse osmosis [RO] water). For alkylglycerol dietary supplementation assay, 10 mM 18-AG stock solution in ethanol was diluted to 100 µM by mixing in the standard medium. For the negative control, the same volume of ethanol was mixed in the medium. The flies were reared in a 25°C incubator under a 12-h light/12-h dark cycle.

### Confocal imaging

For *AGPS* expression analysis, we crossed *AGPS CRIMIC-GAL4* line with *5×UAS-mCD8::GFP* line. Late third instar larvae were fixed on a glass slide by scotch tape and placed on 70°C block for 10 sec. GFP fluorescence was captured using a confocal laser-scanning microscope (SP8, Leica, Germany) equipped with a UPLSAPO 10×/0.40 or a UPLSAPO 40×/0.95 objective lens. Images were analyzed using Leica Application Suite X (LAS X) and ImageJ software^67^.

### Generation of AGPS knockout flies

*AGPS* knockout (*AGPS^KO^*) flies were generated in our laboratory using the CRISPR-Cas9 system. The guide RNA sequence (5’-AATGGCAGCCAAGCGGAATG-3’) was designed using the flyCRISPR target finder (http://targetfinder.flycrispr.neuro.brown.edu/) and cloned into the pU6_3-BbsI-chiRNA vector (Addgene). To achieve homologous recombination, we employed pHD-ScarlessDsRed (Drosophila Genomics Resource Center) as the donor template, which contains 1045 bp upstream and 945 bp downstream of the Cas9-mediated double-strand break site in the *AGPS* genome sequence. Subsequently, the constructed vectors were injected into embryos of vas-cas9 on III (BL #51324). Two distinct *AGPS^KO^* clones, *AGPS^KO^*^1^ and *AGPS^KO2^*, were obtained and outcrossed individually to the control line (*w^1118^*) for five generations. For all experiments, *AGPS^KO1^* was the designated line unless otherwise specified.

### Measurement of pupariation time

To enhance egg production, flies were introduced into vials containing standard food supplemented with a yeast paste for ≥48 h following CO_2_ exposure. Flies were transferred to new vials containing standard food for egg laying for 3 h. These vials were placed inside a plastic bag with water supply to maintain high humidity before measurements. The number of pupae on the vial wall was counted during the light period, 114–192 h AEL. To determine the time when 50% of the animals underwent pupariation (T_50_), we utilized the maximum number of pupae observed in each vial and performed linear regression analysis on the growth curve.

### Quantitative PCR for *Drosophila* larvae and cells

To prepare the whole body sample of late third instar larvae (120 h AEL), ten larvae were collected and rinsed twice in RO water. To isolate the tissues, the larvae were placed in ice-cold phosphate-buffered saline (PBS), and the body wall cuticles were carefully opened using microscissors. After removing the guts and major tracheal organs, the CNS (brain and ventral nerve cord), fat body, and carcass (body wall cuticles) were collected. Tissues were obtained from 5–10 larvae. Whole body or tissue samples were homogenized with a pestle in Sepasol-RNA I Super G (Nacalai Tesque). For cell culture cell, *Drosophila* S2R+ cells were grown in 60-mm dishes at 37°C or 25°C, respectively, until they reached confluence. Then, the cells were washed with PBS and homogenized in Sepasol-RNA I Super G (Nacalai Tesque). Total RNA was extracted following the manufacturer’s protocol.

To isolate *ppk-*positive peripheral sensory neurons from late third instar larvae expressing mCD8 in *ppk*-*GAL4* neurons (*UAS-mCD8::GFP*; *ppk-GAL4*), magnetic bead-based cell sorting was utilized as previously reported^68^, with modifications. Fifty larvae were placed in ice-cold PBS, and the posterior end was excised. The cuticle was inverted, and the internal organs removed. Cuticles were then transferred into a microtube and treated with 50 µg/µL Liberase and 20 mU/µL DNase in 1 mL PBS supplemented with 0.02% Tween 20 (PBSTw) for 30 min at 25°C with agitation. The homogenates were filtered through a 100 µm mesh and further filtered through a 30 µm mesh to remove debris. All apparatuses were coated with 0.1% globulin free-bovine serum albumin (BSA; 016-15111, Wako). Dissociated cells were then collected via centrifugation for 20 min at 300 *× g*, and the supernatant was removed. Next, the cells were resuspended in PBSTw and precleared with protein G Dynabeads for 10 min at 4°C with rotation. The cell suspension was then incubated with α-CD8 antibody-coated protein G Dynabeads for 20 min at 4°C with rotation. Bead-attached cells were collected using a magnetic stand and washed five times with PBS. Total RNA was extracted from bead-bound cells using NucleoSpin RNA Plus XS (Macherey–Nagel) following the manufacturer’s protocol. The RNA concentration was measured using the QuantiFluor RNA System (Promega).

After obtaining the total RNA, cDNA was synthesized using ReverTra Ace. The expression level of the *AGPS* gene was quantified with the ΔΔCt method using *rp49* as a reference gene; KOD SYBR was used for the qPCR, which was performed on StepOne (Applied Biosystems).

### Lipid analysis

For *Drosophila* tissues (10–15 whole bodies and the CNS of late third instar larvae), lipid extraction was performed using the Bligh and Dyer method^69^. Samples were homogenized in MilliQ water on ice and the total lipids were extracted twice using a solvent mixture containing water/methanol/chloroform (1:1:1, v/v). For cultured cells, total lipids were extracted using the methyl-*tert*-butyl ether (MTBE) method^70^. Cells on a dish were detached with PBS and collected in a microtube via centrifugation (1000 × *g*, 5 min). Then, the cells were resuspended in 100 µL MilliQ water, and total lipids were extracted twice using MTBE/methanol/water (10:3:2.5, v/v).

The solvent was evaporated under nitrogen gas, and the final total lipid extract was redissolved in methanol. Phospholipids, including ether-type phospholipids, were analyzed using liquid chromatography with tandem mass spectrometry with LC-30AD coupled to a triple quadrupole mass spectrometer LCMS-8040 (Shimadzu), as previously described^19^. For multiple reaction monitoring (MRM), the transition of [M + H]^+^→ 184.0 was used for PC and [M + H]^+^→[M + H]^+^-141.1 was used for PE. For the product ion scan analysis of ePEs, [M–H]^−^ ions were used as precursor ions derived from PE and scan from m/z:350 to m/z:150.

### Temperature gradient assay

The temperature gradient assay was conducted following the previously described method^36^ with modifications. Aluminum trays [180 mm (W) × 40 mm (D) × 5 mm (H)] were coated with 20 mL of 2% agarose. The left and right ends of the tray were placed on top of two aluminum blocks individually temperature-controlled using a circulating water bath. The surface temperature of the agarose was monitored with a thermometer to create a thermal gradient ranging from 8°C to 35°C (1.5°C/cm).

Late third instar larvae (120 h AEL) were prepared in the same manner as for pupariation measurements. Larvae were collected and washed twice with an 18% sucrose solution and thoroughly rinsed twice with RO water. After a 10-min recovery period, 30–50 larvae were released onto three agarose trays in regions where the surface temperature was approximately 28°C. Each tray was covered with a clear acrylic lid during the assay. The larvae were allowed to explore the agarose under red LED light (> 600 nm) in a black acrylic box. After 10 min, the distribution of larvae on the agarose was captured using a digital camera. To quantify the temperature distribution of larvae, the temperature at which each larva was positioned was calculated using the following equation: Temperature (°C) = Distance from the position at 8°C (cm) × 1.5 (°C/cm) + 8°C. Larvae located within 0.5 cm from the walls or outside the tray were omitted from calculations.

### Thermal two-way choice assay

The assay was conducted on an aluminum tray [90 mm (W) × 130 mm (D) × 8 mm (H)] coated with 25 mL of 2% agarose. The test tray was placed on top of two adjacent aluminum blocks, separated using a thin plastic film (∼1 mm) as spacer. Each block was individually temperature-controlled using a circulating water bath. To prevent agarose from drying, the surface was gently scratched and sprayed with water. The surface temperature on each side of the test tray was confirmed using a thermometer.

At the beginning, 40–60 larvae were released at the border between the two temperature zones and allowed to explore the tray under red LED light in a black acrylic box. After 15 min, the distribution of larvae on the tray was captured using a digital camera, and the number of larvae in each temperature zone tabulated. The avoidance index was calculated using the following formula: (Number of larvae at 20°C − Number of larvae at 29°C)/Total number of larvae on both sides of the test tray. Larvae within the release zone (1 cm wide) were not counted in either temperature zones, and those outside the trays were not included in the calculation.

### Tactile response assays

Late third instar larvae were prepared as for the temperature gradient assay. The gentle-touch assay was performed as described previously^71^. The anterior end of freely moving larvae was lightly touched with a Φ 0.1mm nickel–titanium filament. Larval responses were scored as follows: 0, no response; 1, hesitation; 2, turn or withdrawal of the anterior segments; 3, single reverse contraction; or 4, multiple reverse contraction.

For the mechanical nociception assay, von Frey filaments with different forces were fabricated using a nickel–titanium filament^72^. Filament forces were calibrated by measuring the weight required to bend them and converting the values to force using the following equation: force (mN) = weight (g) × gravity acceleration (9.8 m/s²). Larvae were released on a 60 mm dish under a dissection microscope and stimulated once on the center of their body with a filament per larva. The number of larvae responding to the stimuli within 5 s was recorded.

### Immunohistochemistry

The CNS of late third instar larvae was dissected in a buffer containing 5 mM TES, 10 mM HEPES, 120 mM NaCl, 3 mM KCl, 4 mM MgCl_2_, 2 mM CaCl_2_, 10 mM NaHCO_3_, 10 mM trehalose, 10 mM glucose, and 10 mM sucrose (pH 7.25). The dissected CNS was then placed on 12mmφ cover glass (C012001, Matsunami) coated with poly-L-lysine and fixed with 4% paraformaldehyde and 0.4% Triton X-100 for 30 min at room temperature. Subsequently, samples were washed with PBS containing 0.3% Triton X-100 (PBSTx) and blocked with 5% normal goat serum (NGS) in PBSTx for 30 min at room temperature. Then, samples were incubated with an α-GFP primary antibody (1:500; #A6455, Invitrogen) in 5% NGS/PBSTx overnight at 4°C, followed by washing with PBSTx. Next, samples were incubated with the secondary antibody (α-Rabbit IgG-Alexa488, 1:1000; #A11034, Invitrogen) in PBSTx for 3 h at 4°C and washed with PBSTx. Finally, samples were mounted on a glass slide using Fluoromount (Diagnostic BioSystems). Fluorescence was captured using an Olympus FV1200 IX83 confocal microscope equipped with a 30×/1.05 objective lens.

### S2R+ cell culture, transfection, and generation of stable cell lines

Cells were cultured in Schneider’s *Drosophila* medium (Gibco) supplemented with 10% fetal bovine serum (#10437-028, Gibco) and 50 mg/mL penicillin/50 units/mL streptomycin at 25°C.

For the lipid supplementation experiment, 1 mM lipids were mixed and sonicated in 10 mg/mL fatty acid-free BSA/0.9% NaCl solution for 15 min to form a complex. As negative control, the same volume of solvent (ethanol) was mixed with a BSA/NaCl solution. The lipid-BSA complex was diluted in the culture medium to achieve a final concentration of 100 µM lipids and incubated with the cells for 24 h. The following lipids were purchased: 1-O-hexadecyl-rac-glycerol (16-AG; #sc-205917, Santa Cruz), glycerol 1-oleyl ether (18-AG; #G598710, Tront Research chemicals), and 1-octadecanol (18-OH; #O0006, Tokyo Chemical Industry).

For the functional assay of dTRPA1-A and dTRPA1-B, we established stable cell lines expressing each isoform. dTRPA1-A or dTRPA1-B in pMT vectors were cotransfected with pBS-puro using X-tremeGENE 9 DNA transfection reagent (Roche), and stable cells were selected and maintained in the presence of 10 µg/mL puromycin. Metallothionein promoter-dependent dTRPA1 expression was induced by supplementing cells with 500 µM CuSO_4_ for 24 h. For the functional assay of dTRPA1-A in dAGPS-expressing cells, a cloned cell line stably expressing dAGPS was established. The dAGPS in the pAc5.1-V5-His vector was cotransfected with pBS-puro using the X-tremeGENE 9 DNA transfection reagent. Cells expressing N-term-V5-His-dAGPS were selected in the presence of 10 µg/mL puromycin, and a monoclonal cell line was established using limiting dilution in a 96-well plate (0.5 cell/well). pAc5.1-dTRPA1-A and pAc5.1-EGFP were transiently cotransfected into control or dAGPS stable cells using TransFectin Lipid Reagent (BioRad). After a 24 h incubation, transfection medium was removed and replaced with growth medium, and cells were incubated for another 24 h at 25°C. For the functional assay of dPIEZO, pAc5.1-dPIEZO and pAc5.1-DsRed were transiently cotransfected into control cells using TransFectin Lipid Reagent, as described above.

### Calcium imaging

Cells transiently expressing dPIEZO were loaded with 5 µM Fura 2-AM (Dojindo), 0.02% Pluronic F127, and 500 μM probenecid followed by incubation in standard culture medium for 60 min at 25°C. The same bath solution for electrophysiological experiments was used for measurements. Fura-2 in cells was excited at 340 and 380 nm, and the emission at 510 nm was monitored with a sCMOS camera (Zyla 4.2 Plus, Andor Technology). Time-lapse images were recorded every 3 s. To activate dPIEZO, 10 µM Yoda1 (TOCRIS) was applied by perfusion. The perfusion flow rate was set at 4 mL/min. Ionomycin (5 μM) was applied at the end of the protocol to calibrate the maximum capacity of Ca^2+^ intake. Data were measured and analyzed using the iQ2 software (Andor Technology), and the ratio (340/380) was calculated with Fiji software ^73^.

### Electrophysiology

For whole-cell patch-clamp recordings, the extracellular bath solution contained 140 mM NaCl, 5 mM KCl, 2 mM MgCl_2_, 2 mM CaCl_2_, 10 mM HEPES, and 10 mM glucose (pH 7.4, adjusted with NaOH). The pipette solution contained 140 mM Cs-aspartate, 2LmM MgCl_2_, 0.01 mM CaCl_2_, 1 mM EGTA, and 10 mM HEPES (pH 7.2, adjusted with CsOH). Voltage-clamp recordings were used as previously described^74^, with an Axon 200B (Molecular Devices) amplifier and pCLAMP software (version 10.7; Axon Instruments). The membrane potential was held at −60 mV, and experiments were performed at 25°C. For mechanical stimulation of dPIEZO, a poking stimulation was delivered via membrane indentation (6 μm) using a glass pipette for 150 msec (PCS-5000, Inter Medical). Experiments were performed at 25°C, and the membrane potential was held at −100 mV. The current density was calculated by dividing the maximal current value (nA) by cell capacitance (pF). For heat stimulation of TRPA1, the temperature of the bath solution was initially decreased by perfusing a precooled solution (10°C), then increased by perfusing a prewarmed solution (65°C). The temperature of the solution near the recorded cell was monitored using a TA-29 thermistor (Warner instrument). The temperature threshold for dTRPA1 activation was determined in Arrhenius plots of whole-cell voltage-clamp recordings using Origin software (OriginLab), as previously described^74^. Log scales of currents were plotted against the reciprocal of the absolute temperature (T). The crossing point of the two fitted lines where the slope changed in Arrhenius plots (basal activity vs. post-activation) was defined as the temperature threshold^74^.

### Force curve measurement using high-speed AFM

S2R+ cells, with/without 24-h supplementation with 100 μM 18-AG, were used for analysis. The cells were attached on concanavalin A (Nacalai)-coated cover glass for 30 min at room temperature immediately before the experiment. The force curve measurement was performed in the same bath solution used for electrophysiological experiments but supplemented with 0.01% BSA. The AFM used was a tip-scan high-speed AFM (TS-HS-AFM) combined with an optical microscope^75^. Optical images were used to confirm cell position and the positions of stiffness measurements. The cantilevers used were Olympus AC10 cantilevers (length 9 µm, width 2 µm, and height 0.13 µm), with a carbon tip grown via electron beam deposition (EBD) at its very end. For these measurements, long EBD tips were used as they are reportedly beneficial for cell measurements^76^. Using the thermal sweep method^77^, the cantilevers’ spring constants were determined as 0.093 ± 0.038 Nm^−1^, their resonance frequency as 433 ± 55 kHz, and their Q-factor as 1.34 ± 0.09 (averages and standard deviations of 6 cantilevers). Before measuring the cells, the cantilever’s sensitivity was determined by performing a force map of 50 × 50 curves on a glass substrate and calculating the slope of the force curves using a linear fit. The resulting 2500 slopes were then plotted as a histogram and fitted to a Gaussian curve. The sensitivity was designated as the location of the Gaussian’s maximum. For all measurements, the scan size was 1 × 1 µm, and the pixel resolution set to either 150 × 150 or 200 × 200 pixels.

To measure the mechanical properties of S2R+ cells, the edge of a cell was first located in standard tapping mode with a frame time of 5–10 s. After the cell edge was reached, force curves were recorded with a grid of 50 × 50 curves in between topography lines. This force mapping procedure was based on inline force curve measurements^78^. Simultaneously, optical microscopy images were acquired to assign each force map to a cell position. The frame time during force mapping was 30–35 s. Force curves were recorded with a frequency of 1 kHz. The maximum force was not controlled but was typically <500 pN. Only the flat outer regions of cells were measured because the region close to the nucleus exhibited a steep upward slope that prevented stable scanning conditions. Similarly, it was not possible to scan stably directly on top of the nucleus due to part of the cells fluctuating and being deformed by the small lateral scanning forces.

The acquired force curves were fitted with the Hertz contact mechanics model assuming a tip radius of 5 nm. Force curves were automatically analyzed using home-made software based on IgorPro. Before fitting the force curve with the Hertz model, the linear background was subtracted from the force curves to level the curves.

### Imaging-based membrane property measurement

S2R+ cells, with/without 24 h supplementation of 100 μM 18-AG, were used for analysis. Cells were plated on a non-coat glass bottom dish (35mmφ, No.1S, D11530H, Matsunami) and incubated for 24 h. Subsequently, the cells on the glass were washed with the same bath solution used for electrophysiological experiments and incubated with either 2 µM Flipper-TR (Spirochrome) or 100 nM LipiORDER (Funakoshi) in the bath solution for 15 min at room temperature. For the laurdan imaging, cells were incubated with 2 µM laurdan (Cayman) for 1 hr at room temperature. Images were captured using a Leica TCS SP8 FALCON confocal microscope equipped with a HC PL APO CS2 100×/1.40 oil immersion objective lens, a pulsed supercontinuum white light laser, and HyD-SMD photon counting detectors. Images were acquired at 256 × 256 pixels using 3× zoom. For LipiORDER measurement, cells were excited at 405 nm, and the emission spectrum was measured from 450 to 650 nm with a 25-nm interval. The red to green ratio (R/G ratio) was calculated by dividing fluorescence intensities at 550–650 nm with those at 450–550 nm. For laurdan measurement, the sum of fluorescent intensities at 415–465 nm (I_440_) and 465–515 nm (I_490_) were measured with 405 nm excitation. The GP score was calculated by (I_440_ - I_490_) / (I_440_ + I_490_). For Flipper-TR measurements, we performed excitation at 488 nm and emission at 550–650 nm. The fluorescence lifetimes were calculated from decay curves using Leica LAS X SingleMoleculeDetection software. The fitting model was an n-exponential reconvolution with instrument response function and two exponential components were estimated. The longer decay time was used for subsequent analysis.

### Statistical analysis

Data are presented as means ± SEMs. The times each experiment was performed (n) is indicated either in figure legends or panels. To compare two samples, unpaired, two-tailed Student’s *t* tests were performed. To performed multiple comparison, Tukey’s test or Dunnett’s test was used. For comparison of membrane property assay, nonparametric Mann–Whitney U test (for two samples) and Kruskal–Wallis tests followed by Steel’s tests (for multiple comparisons vs. control) were performed. All statistical analyses were conducted using Prism 10 (GraphPad) or JMP 14.2 software (SAS Institute). A p-value <0.05 was considered statistically significant.

## ACKNOWLEDGMENTS

The fly stocks used in this study were obtained from the Bloomington Drosophila Stock Center, and NIG-FLY Stock Center. We thank Naomi Fukuta, Terumi Hashimoto, and Dr. Jing Lei (ExCELLS/NIPS) for supporting fly maintenance and molecular cloning experiments. We appreciate Dr. Masato Umeda (HoloBio Inc.), Dr. Akifumi Shiomi (RIKEN), and Dr. Akira Murakami (University of Shizuoka) for providing their valuable inputs to the project. This work was supported by Grant-in-aid for Scientific research 19K23790 and 21K15192 (for T. Suito), 21H02531 (for T. Sokabe), and 22H04926 (for T.N.) from Japan Society for the Promotion of Science (JSPS) and Ministry of Education, Culture, Sports, Science and Technology, Grant-in-Aid for Transformative Research Areas ― Platforms for Advanced Technologies and Research Resources “Advanced Bioimaging Support” JP22H04926 (for T. Sokabe) from JSPS, and the Japan Agency for Medical Research and Development (AMED)-PRIME 23gm6510014h0002 (for T. Sokabe) from AMED.

## AUTHORS CONTRIBUTION

T. Suito contributed to designing, conducting, and analyzing most of the experiments and preparing the draft and the final version of the manuscript. T. Sokabe contributed to supervising the project and preparing the draft and the final version of the manuscript. M. Tominaga contributed to supervising the project. K.N. contributed to supervising and performing the LC-MS lipid analysis. X.D. and S.S. contributed to performing fly behavioral experiments. C.G. and T.U. contributed to supervising and conducting the force curve measurement using AFM. M. Tsutsumi and T.N. contributed to supervising chemical probe-based membrane property measurement. Y.H. contributed to dPIEZO cloning and supervising the calcium measurement experiment.

## DECLARATION OF INTERESTS

The authors declare no competing interests.

## Notes

### Competing Interest Statement

The authors have declared no competing interest.

### Summary of Updates

The electrophysiological recording of PIEZO channel added (new Figure 4); Membrane property measurement updated; thermotaxis assay updated; AGPS expression pattern added; lipid profile in larvae updated; ePL precursor supplementation assay added; whole manuscript structure updated.

## REFERENCES

1. Caterina, M.J., Schumacher, M.A., Tominaga, M., Rosen, T.A., Levine, J.D., and Julius, D. (1997). The capsaicin receptor: a heat-activated ion channel in the pain pathway. Nature 389, 816–824. 10.1038/39807.

2. Dhaka, A., Murray, A.N., Mathur, J., Earley, T.J., Petrus, M.J., and Patapoutian, A. (2007). TRPM8 is required for cold sensation in mice. Neuron 54, 371–378. 10.1016/j.neuron.2007.02.024.

3. Moqrich, A., Hwang, S.W., Earley, T.J., Petrus, M.J., Murray, A.N., Spencer, K.S., Andahazy, M., Story, G.M., and Patapoutian, A. (2005). Impaired thermosensation in mice lacking TRPV3, a heat and camphor sensor in the skin. Science 307, 1468–1472. 10.1126/science.1108609.

4. Coste, B., Mathur, J., Schmidt, M., Earley, T.J., Ranade, S., Petrus, M.J., Dubin, A.E., and Patapoutian, A. (2010). Piezo1 and Piezo2 are essential components of distinct mechanically activated cation channels. Science 330, 55–60. 10.1126/science.1193270.

5. Tominaga, M., Caterina, M.J., Malmberg, A.B., Rosen, T.A., Gilbert, H., Skinner, K., Raumann, B.E., Basbaum, A.I., and Julius, D. (1998). The cloned capsaicin receptor integrates multiple pain-producing stimuli. Neuron 21, 531–543. 10.1016/s0896-6273(00)80564-4.

6. Ranade, S.S., Woo, S.H., Dubin, A.E., Moshourab, R.A., Wetzel, C., Petrus, M., Mathur, J., Begay, V., Coste, B., Mainquist, J., et al. (2014). Piezo2 is the major transducer of mechanical forces for touch sensation in mice. Nature 516, 121–125. 10.1038/nature13980.

7. Kashio, M., and Tominaga, M. (2022). TRP channels in thermosensation. Curr Opin Neurobiol 75, 102591. 10.1016/j.conb.2022.102591.

8. Ciardo, M.G., and Ferrer-Montiel, A. (2017). Lipids as central modulators of sensory TRP channels. Biochim Biophys Acta Biomembr 1859, 1615–1628. 10.1016/j.bbamem.2017.04.012.

9. Romero, L.O., Massey, A.E., Mata-Daboin, A.D., Sierra-Valdez, F.J., Chauhan, S.C., Cordero-Morales, J.F., and Vasquez, V. (2019). Dietary fatty acids fine-tune Piezo1 mechanical response. Nat Commun 10, 1200. 10.1038/s41467-019-09055-7.

10. Romero, L.O., Caires, R., Kaitlyn Victor, A., Ramirez, J., Sierra-Valdez, F.J., Walsh, P., Truong, V., Lee, J., Mayor, U., Reiter, L.T., et al. (2023). Linoleic acid improves PIEZO2 dysfunction in a mouse model of Angelman Syndrome. Nat Commun 14, 1167. 10.1038/s41467-023-36818-0.

11. Tsuchiya, M., Hara, Y., Okuda, M., Itoh, K., Nishioka, R., Shiomi, A., Nagao, K., Mori, M., Mori, Y., Ikenouchi, J., et al. (2018). Cell surface flip-flop of phosphatidylserine is critical for PIEZO1-mediated myotube formation. Nat Commun 9, 2049. 10.1038/s41467-018-04436-w.

12. Bradshaw, H.B., Raboune, S., and Hollis, J.L. (2013). Opportunistic activation of TRP receptors by endogenous lipids: exploiting lipidomics to understand TRP receptor cellular communication. Life Sci 92, 404–409. 10.1016/j.lfs.2012.11.008.

13. Sisignano, M., Bennett, D.L., Geisslinger, G., and Scholich, K. (2014). TRP-channels as key integrators of lipid pathways in nociceptive neurons. Prog Lipid Res 53, 93–107. 10.1016/j.plipres.2013.11.002.

14. Taberner, F.J., Fernandez-Ballester, G., Fernandez-Carvajal, A., and Ferrer-Montiel, A. (2015). TRP channels interaction with lipids and its implications in disease. Biochim Biophys Acta 1848, 1818–1827. 10.1016/j.bbamem.2015.03.022.

15. Cordero-Morales, J.F., and Vasquez, V. (2018). How lipids contribute to ion channel function, a fat perspective on direct and indirect interactions. Curr Opin Struct Biol 51, 92–98. 10.1016/j.sbi.2018.03.015.

16. Bazinet, R.P., and Laye, S. (2014). Polyunsaturated fatty acids and their metabolites in brain function and disease. Nat Rev Neurosci 15, 771–785. 10.1038/nrn3820.

17. Custers, Emma, E.M., Kiliaan, and Amanda, J. (2022). Dietary lipids from body to brain. Prog Lipid Res 85, 101144. 10.1016/j.plipres.2021.101144.

18. Vasquez, V., Krieg, M., Lockhead, D., and Goodman, M.B. (2014). Phospholipids that contain polyunsaturated fatty acids enhance neuronal cell mechanics and touch sensation. Cell Rep 6, 70–80. 10.1016/j.celrep.2013.12.012.

19. Suito, T., Nagao, K., Takeuchi, K., Juni, N., Hara, Y., and Umeda, M. (2020). Functional expression of Delta12 fatty acid desaturase modulates thermoregulatory behaviour in Drosophila. Sci Rep 10, 11798. 10.1038/s41598-020-68601-2.

20. Snyder, F. (1999). The ether lipid trail: a historical perspective. Biochim Biophys Acta 1436, 265–278. 10.1016/s0005-2760(98)00172-6.

21. Braverman, N.E., and Moser, A.B. (2012). Functions of plasmalogen lipids in health and disease. Biochim Biophys Acta 1822, 1442–1452. 10.1016/j.bbadis.2012.05.008.

22. Dorninger, F., Forss-Petter, S., and Berger, J. (2017). From peroxisomal disorders to common neurodegenerative diseases - the role of ether phospholipids in the nervous system. FEBS Lett 591, 2761–2788. 10.1002/1873-3468.12788.

23. Rudiger, M., Kolleck, I., Putz, G., Wauer, R.R., Stevens, P., and Rustow, B. (1998). Plasmalogens effectively reduce the surface tension of surfactant-like phospholipid mixtures. Am J Physiol 274, L143–148. 10.1152/ajplung.1998.274.1.L143.

24. Flasinski, M., Wydro, P., Hac-Wydro, K., and Dynarowicz-Latka, P. (2013). Cholesterol as a factor regulating the influence of natural (PAF and lysoPAF) vs synthetic (ED) ether lipids on model lipid membranes. Biochim Biophys Acta 1828, 2700–2708. 10.1016/j.bbamem.2013.07.024.

25. Ford, D.A., and Hale, C.C. (1996). Plasmalogen and anionic phospholipid dependence of the cardiac sarcolemmal sodium-calcium exchanger. FEBS Lett 394, 99–102. 10.1016/0014-5793(96)00930-1.

26. A. Razeto, A., Mattiroli, F., Carpanelli, E., Aliverti, A., Pandini, V., Coda, A., and Mattevi, (2007). The Crucial Step in Ether Phospholipid Biosynthesis: Structural Basis of a Noncanonical Reaction Associated with a Peroxisomal Disorder. Structure 15, 683–692. 10.1016/j.str.2007.04.009.

27. Im, S.H., and Galko, M.J. (2012). Pokes, sunburn, and hot sauce: Drosophila as an emerging model for the biology of nociception. Dev Dyn 241, 16–26. 10.1002/dvdy.22737.

28. Shiomi, A., Nagao, K., Yokota, N., Tsuchiya, M., Kato, U., Juni, N., Hara, Y., Mori, M.X., Mori, Y., Ui-Tei, K., et al. (2021). Extreme deformability of insect cell membranes is governed by phospholipid scrambling. Cell Rep 35, 109219. 10.1016/j.celrep.2021.109219.

29. Fitzner, D., Bader, J.M., Penkert, H., Bergner, C.G., Su, M., Weil, M.T., Surma, M.A., Mann, M., Klose, C., and Simons, M. (2020). Cell-Type- and Brain-Region-Resolved Mouse Brain Lipidome. Cell Rep 32, 108132. 10.1016/j.celrep.2020.108132.

30. Kim, S.E., Coste, B., Chadha, A., Cook, B., and Patapoutian, A. (2012). The role of Drosophila Piezo in mechanical nociception. Nature 483, 209–212. 10.1038/nature10801.

31. Zhong, L., Bellemer, A., Yan, H., Ken, H., Jessica, R., Hwang, R.Y., Pitt, G.S., and Tracey, W.D. (2012). Thermosensory and nonthermosensory isoforms of Drosophila melanogaster TRPA1 reveal heat-sensor domains of a thermoTRP Channel. Cell Rep 1, 43–55. 10.1016/j.celrep.2011.11.002.

32. Gong, J., Chen, J., Gu, P., Shang, Y., Ruppell, K.T., Yang, Y., Wang, F., Wen, Q., and Xiang, Y. (2022). Shear stress activates nociceptors to drive Drosophila mechanical nociception. Neuron 110, 3727–3742 e3728. 10.1016/j.neuron.2022.08.015.

33. Zhong, L., Hwang, R.Y., and Tracey, W.D. (2010). Pickpocket is a DEG/ENaC protein required for mechanical nociception in Drosophila larvae. Curr Biol 20, 429–434. 10.1016/j.cub.2009.12.057.

34. Tracey, W.D., Jr., Wilson, R.I., Laurent, G., and Benzer, S. (2003). painless, a Drosophila gene essential for nociception. Cell 113, 261–273. 10.1016/s0092-8674(03)00272-1.

35. Luo, J., Shen, W.L., and Montell, C. (2017). TRPA1 mediates sensation of the rate of temperature change in Drosophila larvae. Nature neuroscience 20, 34–41. 10.1038/nn.4416.

36. Sokabe, T., Chen, H.C., Luo, J., and Montell, C. (2016). A Switch in Thermal Preference in Drosophila Larvae Depends on Multiple Rhodopsins. Cell Rep 17, 336–344. 10.1016/j.celrep.2016.09.028.

37. Gu, P., Gong, J., Shang, Y., Wang, F., Ruppell, K.T., Ma, Z., Sheehan, A.E., Freeman, M.R., and Xiang, Y. (2019). Polymodal Nociception in Drosophila Requires Alternative Splicing of TrpA1. Curr Biol 29, 3961–3973 e3966. 10.1016/j.cub.2019.09.070.

38. Rosenzweig, M., Kang, K., and Garrity, P.A. (2008). Distinct TRP channels are required for warm and cool avoidance in Drosophila melanogaster. Proc Natl Acad Sci U S A 105, 14668–14673. 10.1073/pnas.0805041105.

39. Kwon, Y., Shen, W.L., Shim, H.S., and Montell, C. (2010). Fine thermotactic discrimination between the optimal and slightly cooler temperatures via a TRPV channel in chordotonal neurons. J Neurosci 30, 10465–10471. 10.1523/JNEUROSCI.1631-10.2010.

40. Brites, P., Ferreira, A.S., da Silva, T.F., Sousa, V.F., Malheiro, A.R., Duran, M., Waterham, H.R., Baes, M., and Wanders, R.J. (2011). Alkyl-glycerol rescues plasmalogen levels and pathology of ether-phospholipid deficient mice. PLoS One 6, e28539. 10.1371/journal.pone.0028539.

41. Ziegler, A.B., Thiele, C., Tenedini, F., Richard, M., Leyendecker, P., Hoermann, A., Soba, P., and Tavosanis, G. (2017). Cell-Autonomous Control of Neuronal Dendrite Expansion via the Fatty Acid Synthesis Regulator SREBP. Cell Rep 21, 3346–3353. 10.1016/j.celrep.2017.11.069.

42. Meltzer, S., Bagley, J.A., Perez, G.L., O’Brien, C.E., DeVault, L., Guo, Y., Jan, L.Y., and Jan, Y.N. (2017). Phospholipid Homeostasis Regulates Dendrite Morphogenesis in Drosophila Sensory Neurons. Cell Rep 21, 859–866. 10.1016/j.celrep.2017.09.089.

43. Syeda, R., Xu, J., Dubin, A.E., Coste, B., Mathur, J., Huynh, T., Matzen, J., Lao, J., Tully, D.C., Engels, I.H., et al. (2015). Chemical activation of the mechanotransduction channel Piezo1. Elife 4. 10.7554/eLife.07369.

44. Caires, R., Sierra-Valdez, F.J., Millet, J.R.M., Herwig, J.D., Roan, E., Vasquez, V., and Cordero-Morales, J.F. (2017). Omega-3 Fatty Acids Modulate TRPV4 Function through Plasma Membrane Remodeling. Cell Rep 21, 246–258. 10.1016/j.celrep.2017.09.029.

45. Colom, A., Derivery, E., Soleimanpour, S., Tomba, C., Molin, M.D., Sakai, N., Gonzalez-Gaitan, M., Matile, S., and Roux, A. (2018). A fluorescent membrane tension probe. Nat Chem 10, 1118–1125. 10.1038/s41557-018-0127-3.

46. Valanciunaite, J., Kempf, E., Seki, H., Danylchuk, D.I., Peyrieras, N., Niko, Y., and Klymchenko, A.S. (2020). Polarity Mapping of Cells and Embryos by Improved Fluorescent Solvatochromic Pyrene Probe. Anal Chem 92, 6512–6520. 10.1021/acs.analchem.0c00023.

47. Parasassi, T., De Stasio, G., Ravagnan, G., Rusch, R.M., and Gratton, E. (1991). Quantitation of lipid phases in phospholipid vesicles by the generalized polarization of Laurdan fluorescence. Biophys J 60, 179–189. 10.1016/S0006-3495(91)82041-0.

48. Zou, Y., Henry, W.S., Ricq, E.L., Graham, E.T., Phadnis, V.V., Maretich, P., Paradkar, S., Boehnke, N., Deik, A.A., Reinhardt, F., et al. (2020). Plasticity of ether lipids promotes ferroptosis susceptibility and evasion. Nature 585, 603–608. 10.1038/s41586-020-2732-8.

49. Astudillo, A.M., Balboa, M.A., and Balsinde, J. (2023). Compartmentalized regulation of lipid signaling in oxidative stress and inflammation: Plasmalogens, oxidized lipids and ferroptosis as new paradigms of bioactive lipid research. Prog Lipid Res 89, 101207. 10.1016/j.plipres.2022.101207.

50. Hirata, Y., Cai, R., Volchuk, A., Steinberg, B.E., Saito, Y., Matsuzawa, A., Grinstein, S., and Freeman, S.A. (2023). Lipid peroxidation increases membrane tension, Piezo1 gating, and cation permeability to execute ferroptosis. Curr Biol 33, 1282–1294 e1285. 10.1016/j.cub.2023.02.060.

51. Morfin, N., Fillier, T.A., Pham, T.H., Goodwin, P.H., Thomas, R.H., and Guzman-Novoa, E. (2022). First insights into the honey bee (Apis mellifera) brain lipidome and its neonicotinoid-induced alterations associated with reduced self-grooming behavior. J Adv Res 37, 75–89. 10.1016/j.jare.2021.08.007.

52. Wynalda, K.M., and Murphy, R.C. (2010). Low-concentration ozone reacts with plasmalogen glycerophosphoethanolamine lipids in lung surfactant. Chem Res Toxicol 23, 108–117. 10.1021/tx900306p.

53. Gallego-Garcia, A., Monera-Girona, A.J., Pajares-Martinez, E., Bastida-Martinez, E., Perez-Castano, R., Iniesta, A.A., Fontes, M., Padmanabhan, S., and Elias-Arnanz, M. (2019). A bacterial light response reveals an orphan desaturase for human plasmalogen synthesis. Science 366, 128–132. 10.1126/science.aay1436.

54. Werner, E.R., Keller, M.A., Sailer, S., Lackner, K., Koch, J., Hermann, M., Coassin, S., Golderer, G., Werner-Felmayer, G., Zoeller, R.A., et al. (2020). The TMEM189 gene encodes plasmanylethanolamine desaturase which introduces the characteristic vinyl ether double bond into plasmalogens. Proc Natl Acad Sci U S A 117, 7792–7798. 10.1073/pnas.1917461117.

55. Chintapalli, V.R., Wang, J., and Dow, J.A.T. (2007). Using FlyAtlas to identify better Drosophila melanogaster models of human disease. Nature Genetics 39, 715–720. 10.1038/ng2049.

56. Yang, X., Lin, C., Chen, X., Li, S., Li, X., and Xiao, B. (2022). Structure deformation and curvature sensing of PIEZO1 in lipid membranes. Nature 604, 377–383. 10.1038/s41586-022-04574-8.

57. Hamada, T., Kishimoto, Y., Nagasaki, T., and Takagi, M. (2011). Lateral phase separation in tense membranes. Soft Matter 7. 10.1039/c1sm05948c.

58. Oglecka, K., Rangamani, P., Liedberg, B., Kraut, R.S., and Parikh, A.N. (2014). Oscillatory phase separation in giant lipid vesicles induced by transmembrane osmotic differentials. Elife 3, e03695. 10.7554/eLife.03695.

59. Chen, D., and Santore, M.M. (2014). Three dimensional (temperature-tension-composition) phase map of mixed DOPC-DPPC vesicles: Two solid phases and a fluid phase coexist on three intersecting planes. Biochim Biophys Acta 1838, 2788–2797. 10.1016/j.bbamem.2014.07.014.

60. Guo, Y.R., and MacKinnon, R. (2017). Structure-based membrane dome mechanism for Piezo mechanosensitivity. Elife 6. 10.7554/eLife.33660.

61. Lewis, A.H., and Grandl, J. (2015). Mechanical sensitivity of Piezo1 ion channels can be tuned by cellular membrane tension. Elife 4. 10.7554/eLife.12088.

62. Zhang, M., Ma, Y., Ye, X., Zhang, N., Pan, L., and Wang, B. (2023). TRP (transient receptor potential) ion channel family: structures, biological functions and therapeutic interventions for diseases. Signal Transduct Target Ther 8, 261. 10.1038/s41392-023-01464-x.

63. Delmas, P., Parpaite, T., and Coste, B. (2022). PIEZO channels and newcomers in the mammalian mechanosensitive ion channel family. Neuron 110, 2713–2727. 10.1016/j.neuron.2022.07.001.

64. Barnes-Velez, J.A., Aksoy Yasar, F.B., and Hu, J. (2023). Myelin lipid metabolism and its role in myelination and myelin maintenance. Innovation (Camb) 4, 100360. 10.1016/j.xinn.2022.100360.

65. Nishimoto, R., Derouiche, S., Eto, K., Deveci, A., Kashio, M., Kimori, Y., Matsuoka, Y., Morimatsu, H., Nabekura, J., and Tominaga, M. (2021). Thermosensitive TRPV4 channels mediate temperature-dependent microglia movement. Proc Natl Acad Sci U S A 118. 10.1073/pnas.2012894118.

66. Acheta, J., Bhatia, U., Haley, J., Hong, J., Rich, K., Close, R., Bechler, M.E., Belin, S., and Poitelon, Y. (2022). Piezo channels contribute to the regulation of myelination in Schwann cells. Glia 70, 2276–2289. 10.1002/glia.24251.

67. Schneider, C.A., Rasband, W.S., and Eliceiri, K.W. (2012). NIH Image to ImageJ: 25 years of image analysis. Nat Methods 9, 671–675. 10.1038/nmeth.2089.

68. Turner, H.N., Armengol, K., Patel, A.A., Himmel, N.J., Sullivan, L., Iyer, S.C., Bhattacharya, S., Iyer, E.P.R., Landry, C., Galko, M.J., and Cox, D.N. (2016). The TRP Channels Pkd2, NompC, and Trpm Act in Cold-Sensing Neurons to Mediate Unique Aversive Behaviors to Noxious Cold in Drosophila. Curr Biol 26, 3116–3128. 10.1016/j.cub.2016.09.038.

69. Bligh, E.G., and Dyer, W.J. (1959). A rapid method of total lipid extraction and purification. Can J Biochem Physiol 37, 911–917. 10.1139/o59-099.

70. Matyash, V., Liebisch, G., Kurzchalia, T.V., Shevchenko, A., and Schwudke, D. (2008). Lipid extraction by methyl-tert-butyl ether for high-throughput lipidomics. J Lipid Res 49, 1137–1146. 10.1194/jlr.D700041-JLR200.

71. Kernan, M., Cowan, D., and Zuker, C. (1994). Genetic dissection of mechanosensory transduction: mechanoreception-defective mutations of Drosophila. Neuron 12, 1195–1206. 10.1016/0896-6273(94)90437-5.

72. Lopez-Bellido, R., and Galko, M.J. (2020). An Improved Assay and Tools for Measuring Mechanical Nociception in Drosophila Larvae. J Vis Exp. 10.3791/61911.

73. Schindelin, J., Arganda-Carreras, I., Frise, E., Kaynig, V., Longair, M., Pietzsch, T., Preibisch, S., Rueden, C., Saalfeld, S., Schmid, B., et al. (2012). Fiji: an open-source platform for biological-image analysis. Nat Methods 9, 676-682. 10.1038/nmeth.2019.

74. Nguyen, T.H.D., Chapman, S., Kashio, M., Saito, C., Strom, T., Yasui, M., and Tominaga, M. (2022). Single amino acids set apparent temperature thresholds for heat-evoked activation of mosquito transient receptor potential channel TRPA1. J Biol Chem 298, 102271. 10.1016/j.jbc.2022.102271.

75. Fukuda, S., Uchihashi, T., Iino, R., Okazaki, Y., Yoshida, M., Igarashi, K., and Ando, T. (2013). High-speed atomic force microscope combined with single-molecule fluorescence microscope. Rev Sci Instrum 84, 073706. 10.1063/1.4813280.

76. Shibata, M., Uchihashi, T., Ando, T., and Yasuda, R. (2015). Long-tip high-speed atomic force microscopy for nanometer-scale imaging in live cells. Sci Rep 5, 8724. 10.1038/srep08724.

77. Hutter, J.L., and Bechhoefer, J. (1993). Calibration of atomic-force microscope tips. Review of Scientific Instruments 64, 1868–1873. 10.1063/1.1143970.

78. Ganser, C., and Uchihashi, T. (2018). Microtubule self-healing and defect creation investigated by in-line force measurements during high-speed atomic force microscopy imaging. Nanoscale 11, 125–135. 10.1039/c8nr07392a.

