## Supplementary figures for "Ether phospholipids modulate somatosensory responses by tuning multiple receptor functions in *Drosophila*"

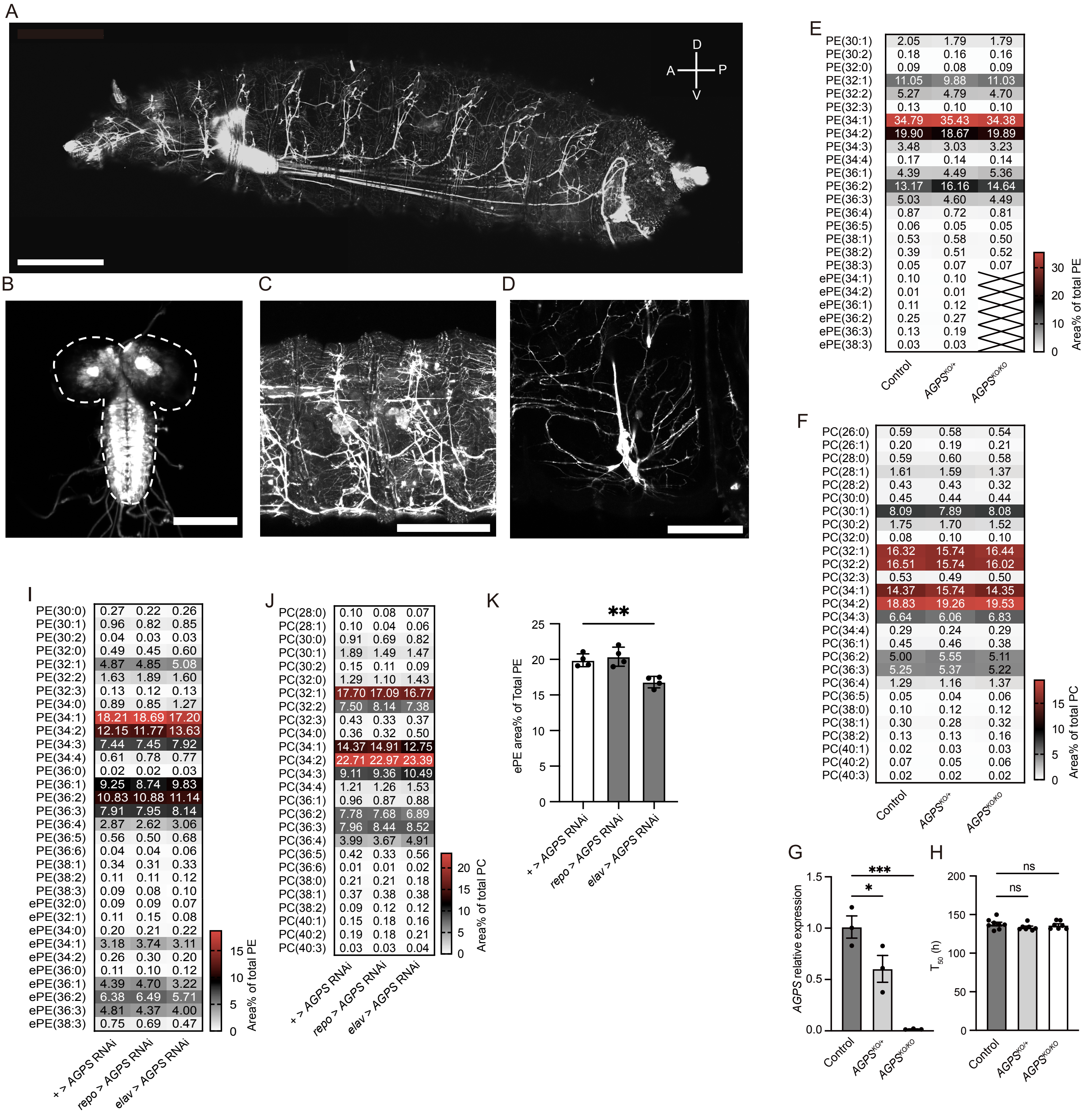

**Figure S1.** ***AGPS*** **expression pattern and lipid profile in** ***AGPS*** **KO third instar larva. Related to Figure 1.**

**A-D** Images of *AGPS* expression patterns at late third instar stage in whole body (**A**, Scale bar: 500 μm), central nervous system (**B**, Scale bar: 200 μm), peripheral neurons (**C**, Scale bar: 200 μm), and class IV multidendritic neurons (**D**, Scale bar: 100 μm). GFP was expressed using *AGPS-CRIMIC-GAL4*. **E, F** Colormap indicating the lipid composition of PE (**E**) and PC (**F**) species (area% of total PE or PC) in the whole body of $w^{1118}$ (control), heterozygous *AGPS* knockout ($AGPS^{KO/+}$), and homozygous *AGPS* knockout ($AGPS^{KO/KO}$) third instar larvae (n = 4, each). Phospholipid molecules are listed as PE (X:Y) or PC (X:Y), where X and Y denotes the total number of acyl chains and the total of double bonds in acyl chains, respectively. Cross marks indicate phospholipid species under detectable levels. **G** Relative *AGPS* expression level in the whole body of control ($w^{1118}$), $AGPS^{KO/+}$, and $AGPS^{KO/KO}$ third instar larvae (n = 3). **H** Pupariation time ($T_{50}$) of control (n = 8), $AGPS^{KO/+}$ (n = 7), and $AGPS^{KO/KO}$ (n = 7). Each point represents a biological replicate. Data are presented as mean ± SEM. *p < 0.05; *** p < 0.001, Dunnett test (vs. control). **I, J** Colormaps of the lipid composition for PE (**I**) and PC (**J**) species (area% of total PE or PC) in the CNS of control (+ > *AGPS* RNAi), pan-glial *AGPS* knockdown (*repo* > *AGPS* RNAi), Pan-neural *AGPS* knockdown (*elav* > *AGPS* RNAi) in third instar larvae (n = 4, each). **K** Total ePE area (% of total PE in the CNS) of third instar larvae (n = 4). **p < 0.01; Dunnett's test.

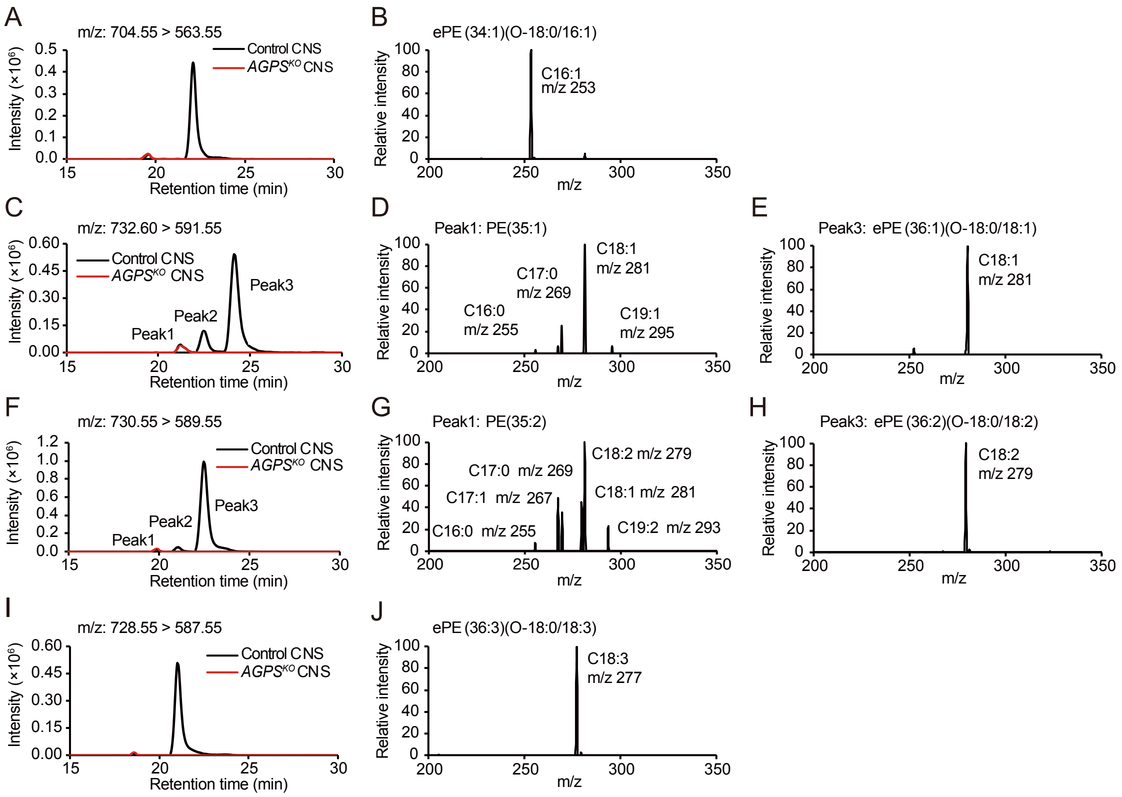


**Figure S2. Product ion scan analysis of major ePEs in *Drosophila* third instar larvae. Related to Figure 1.**

**A** Chromatogram of MRM transition at m/z: 704.55 >563.55 in the CNS of control (black) and $AGPS^{KO}$ (red) larvae. **B** Product ion scan analysis of the peak in the control CNS (**A**). A fragment coinciding with C16:1 considered as derived from the *sn-2* ester bond of ePE (34:1) (O-18:0/16:1). **C** Chromatogram of MRM transition at m/z: 732.60 > 591.55 in control and $AGPS^{KO}$ CNS. The peak2 signal is derived from the isotope of ePE (36:2). **D** Product ion scan analysis of peak1 in the control CNS (**C**). Fragments estimated to be derived from odd-chain fatty acid containing PE (35:1). **E** Product ion scan analysis of peak3 in the control CNS (**C**). A fragment coinciding with C18:1 considered as derived from the *sn-2* ester bond of ePE (36:1) (O-18:0/18:1). **F** Chromatogram of MRM transition at m/z: 730.55 >589.55 in control and $AGPS^{KO}$ CNS. The peak2 signal is derived from the isotope of ePE (36:3). **G** Product ion scan analysis of peak1 in the control CNS (**F**). Fragments estimated to be derived from odd-chain fatty acid containing PE (35:2). **H** Product ion scan analysis of peak3 in the control CNS (**F**). A fragment coinciding with C18:2 estimated to be derived from the *sn-2* ester bond of ePE (36:2) (O-18:0/18:2). **I** Chromatogram of MRM transition at m/z: 728.55 >587.55 in control and $AGPS^{KO}$ CNS. **J** Product ion scan analysis of the peak in the control CNS (**I**). A fragment coinciding with C18:3 considered as derived from the *sn-2* ester bond of ePE (36:3) (O-18:0/18:3).

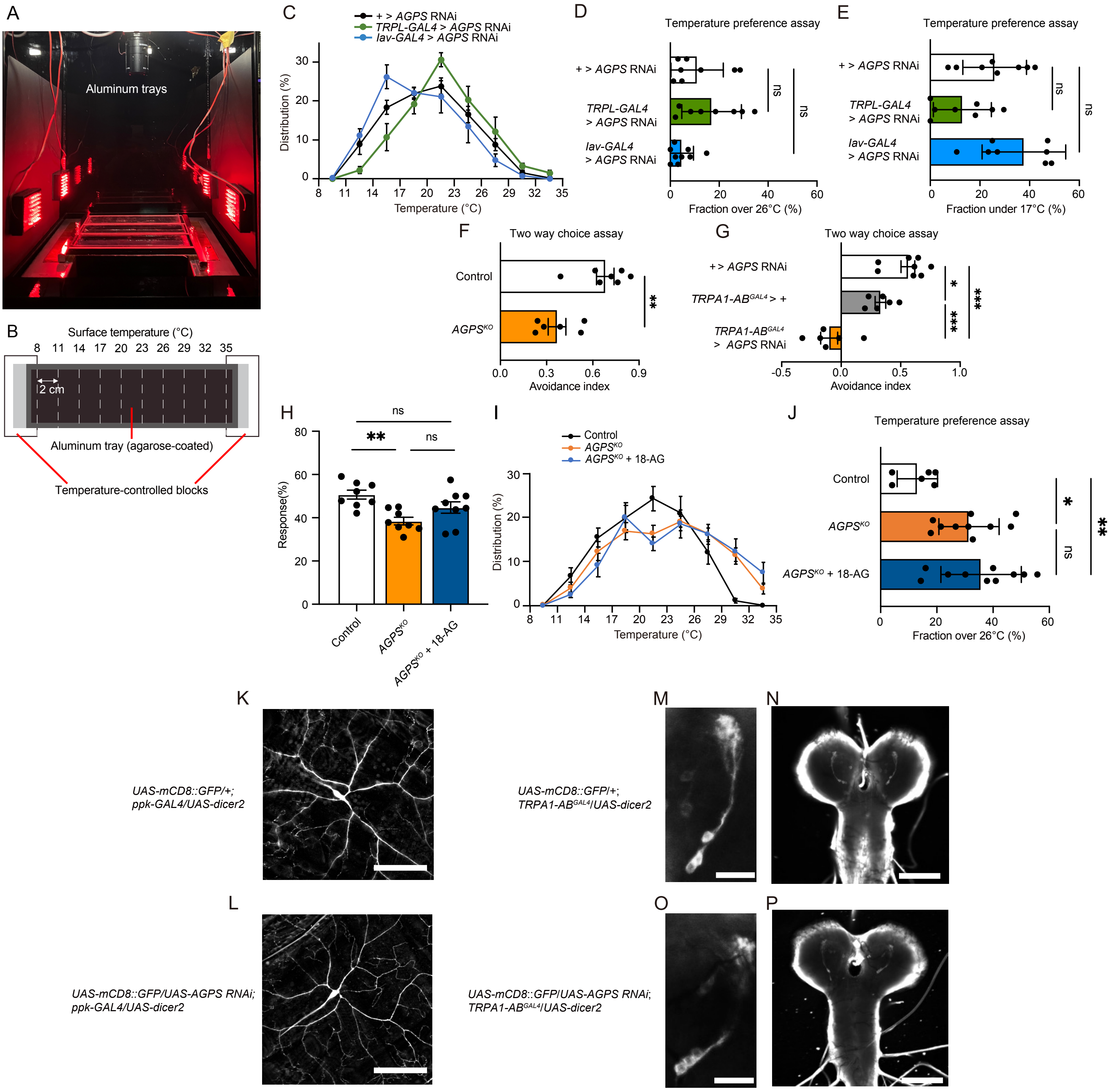


**Figure S3. The phenotypes in *AGPS* knocked-down larvae or rescue of behavioral defect in AGPS KO. Related to Figure 2.**

**A, B** The image and cartoon of thermal gradient setup. **C, D, E** Temperature preference assay for late third instar larvae of control (*UAS-AGPS* RNAi/+, n = 8), and *AGPS* knockdown by *TRPL-GAL4* (*UAS-dicer-2/UAS-AGPS* RNAi;*TRPL-GAL4*/+, n = 8) and *Iav-GAL4* (*UAS-dicer-2/UAS-AGPS* RNAi;*Iav-GAL4*/+, n = 9). the fraction of larvae distributed over 26ºC zones (**D**) or under 17ºC zones (**E**). no significant difference was observed using Dunnett's test. **F, G** Avoidance index in the thermal two-way choice assay at 29ºC vs. 20ºC of $w^{1118}$ (control, n = 7) and $AGPS^{KO}$ (n = 6) **(F)**, or controls ($TRPA1\text{-}AB^{GAL4}$/+, n = 6 and *AGPS* RNAi/+, n = 8) and *AGPS* knockdown by $TRPA1\text{-}AB^{GAL4}$ (*UAS-dicer-2/UAS-AGPS* RNAi;$TRPA1\text{-}AB^{GAL4}$/+, n = 6) **(G)**. **H** Responses to von Frey filaments (20.1mN) in $w^{1118}$ (control, n = 8) $AGPS^{KO}$ late third instar larvae (n = 8), and $AGPS^{KO}$ supplemented with 100 µM octadecanoyl glycerol ($AGPS^{KO}$ + 18-AG, n = 9). **I, J** Temperature preference assay for late third instar larvae of $w^{1118}$ (control, n = 6), $AGPS^{KO}$ (n = 10), and $AGPS^{KO}$ supplemented with 100 mM octadecanoyl glycerol ($AGPS^{KO}$ + 18-AG, n = 10). Distribution of third instar larvae on a thermal gradient of 8ºC–35ºC (**A**) and the fraction of larvae distributed over 26ºC zones (**J**). Each point represents a biological replicate (**D-H**, **J**). Data are presented as mean ± SEM (**C-J**). ns: no significant difference, *p < 0.05, **p < 0.01, *** p < 0.001; Dunnett's test (**D**, **E**), Student's t-test (**F**) or Tukey's test (**G, H, J**). **K**, **L** Fluorescent images of ppk-positive neurons of late third instar larvae of control (*UAS-mCD8::GFP/+;ppk-GAL4/UAS-dicer2*) (**K**) and *AGPS* knockdown (*UAS-mCD8::GFP/UAS-AGPS* RNAi;*ppk-GAL4/UAS-dicer2*) (**L**). Scale bars, 100 µm. **M-P** Immunohistochemistry images of TRPA1-AB positive neurons in the brain of late third instar larvae of control (*UAS-mCD8::GFP/+*;$TRPA1\text{-}AB^{GAL4}$/*UAS-dicer2*) (**M, N**) and *AGPS* knockdown (*UAS-mCD8::GFP/UAS-AGPS* RNAi;$TRPA1\text{-}AB^{GAL4}$/*UAS-dicer2*) (**O**, **P**). Scale bars, 20 µm (**M**, **O**) or 100 µm (**N**, **P**). **M** and **O** represent magnified images of brain lateral posterior neurons in the CNS in **N** and **P**, respectively. Three or more biological samples were investigated in each genotypes.

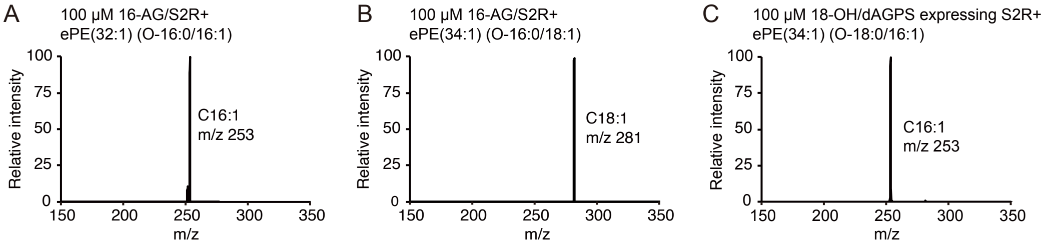


**Figure S4. Product ion scan analysis of major ePEs in ePL-producing S2R+ cells. Related to Figure 3.**

**A** Product ion scan analysis of ePE (32:1) (m/z:674.5) in control S2R+ cells supplemented with 100 μM 16-AG. A fragment coinciding with C16:1 estimated to be derived from the *sn-2* ester bond of ePE (32:1) (O-16:0/16:1). **B** Product ion scan analysis of ePE (34:1) (m/z: 702.55) in control S2R+ cells supplemented with 100 μM 16-AG. A fragment coinciding with C18:1 estimated to be derived from the *sn-2* ester bond of ePE (34:1) (O-16:0/18:1). **C** Product ion scan analysis of ePE (34:1) (m/z: 702.55) in control S2R+ cells supplemented with 100 μM 18-OH. A fragment coinciding with C16:1 estimated to be derived from the *sn-2* ester bond of ePE (34:1) (O-18:0/16:1).

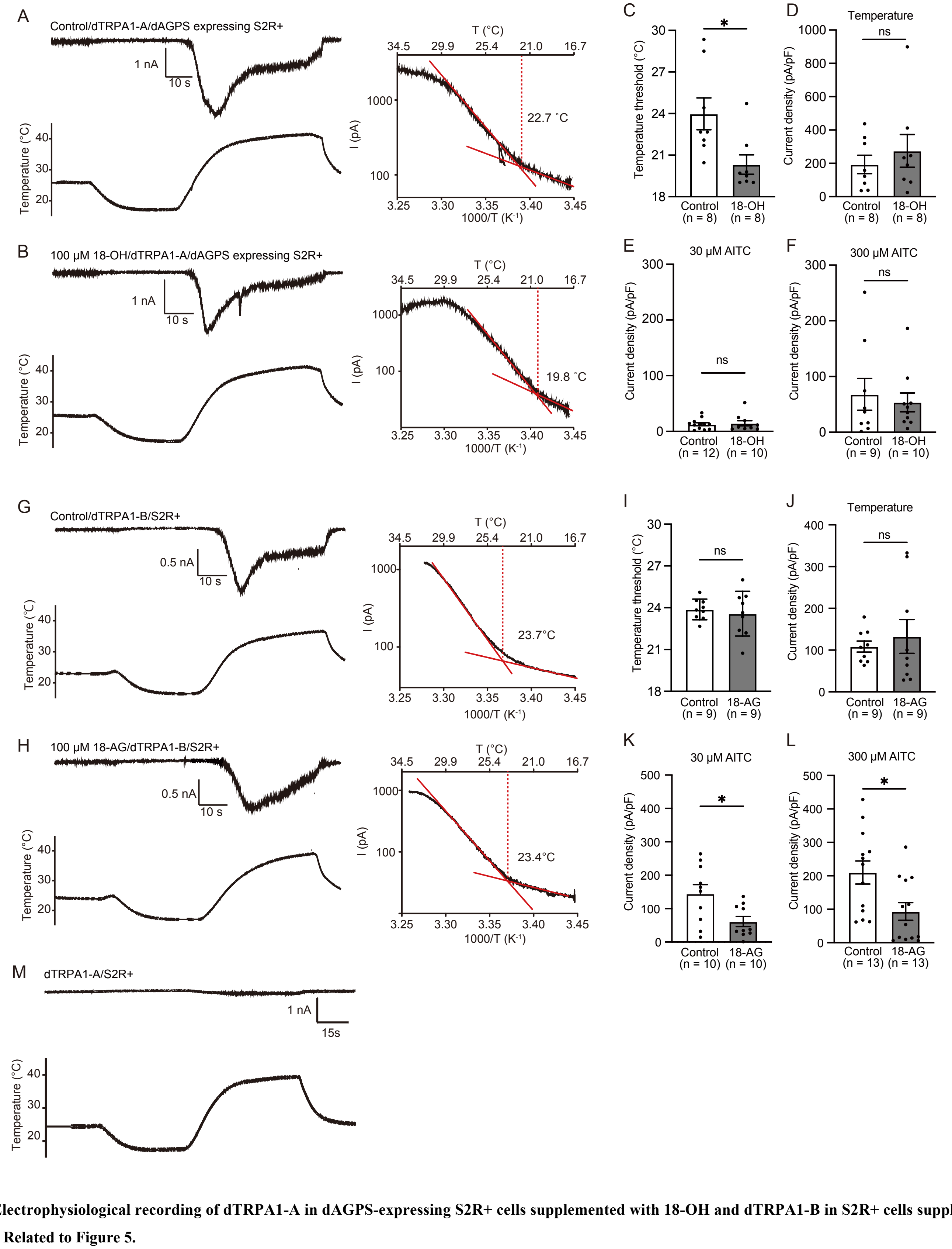


**Figure S5. Electrophysiological recording of dTRPA1-A in dAGPS-expressing S2R+ cells supplemented with 18-OH and dTRPA1-B in S2R+ cells supplemented with 18-AG. Related to Figure 5.**

**A, B** Whole-cell patch-clamp recording of dTRPA1-A activation with temperature stimulation in dAGPS-expressing S2R+ cells (**A**) and dAGPS-expressing S2R+ cells supplemented with 100 μM 18-OH (**B**). Left, typical traces of the recording; right, Arrhenius plots from the traces in the left. Temperature thresholds determined as a crossing point of two fitted lines. **C–F** Quantification of the temperature threshold (**C**) and peak current densities with heat (**D**), 30 μM AITC (**E**), and 300 μM AITC (**F**) stimulation in dAGPS-expressing cells (control) and dAGPS-expressing cells supplemented with 100 μM 18-OH (18-OH). **G, H** Whole-cell patch-clamp recording of dTRPA1-B activation with temperature stimulation in S2R+ cells (**G**) and S2R+ cells supplemented with 100 μM 18-AG (**H**). Left, typical traces of the recording; right, Arrhenius plots from the traces in the left. Temperature thresholds were determined as a crossing point of two fitted lines. **I–L** Quantification of the temperature threshold (**I**) and peak current densities with heat (**J**), 30 μM AITC (**K**), and 300 μM AITC (**L**) stimulation in S2R+ cells (Control) and S2R+ cells supplemented with 100 μM 18-AG (18-AG). Each point represents a biological replicate. The number of the replicates (n) is shown in the panels. Data are presented as mean ± SEM. *$p < 0.05$; Student's t-test. **M** Whole-cell patch-clamp recording of mock transfected S2R+ cells with temperature stimulation.

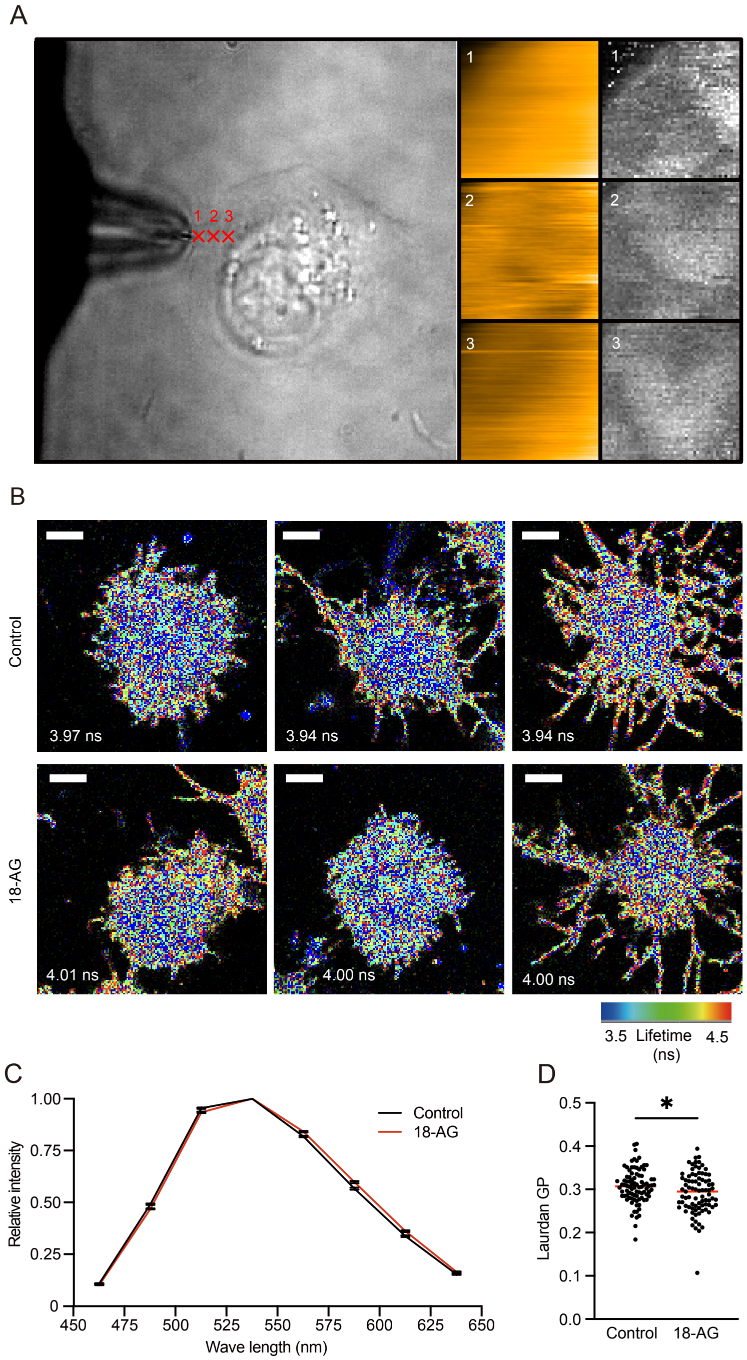


**Figure S6. Representative measurements of physicochemical properties of *Drosophila* cells. Related to Figure 6.**

**A** Representative images of membrane tension measurements in S2R+ cell using AFM. Left, optical image of a target cell and a cantilever (26 × 26 µm). Center, topology imaging (1 × 1 µm) at the points indicated in the left panel. Right, imaging of Young's modulus at the points indicated in the left panel. **B** Representative images of Flipper-TR fluorescent lifetime measurement in S2R+ cells (control, upper panels) and cells supplemented with 100 µM 18-AG (18-AG, lower panel). Scale bars represent 5 µm. Values correspond to the average fluorescent lifetime of the cell membrane. **C** Fluorescence emission spectrum between 450 and 650 nm with a 25-nm interval in LipiORDER analysis for control (black, n = 46) and 100 µM 18-AG supplemented S2R+ cells (red, n = 45). **D** The calculated Laurdan GP score. Each value indicates the average GP score of a single cell. Red horizontal lines represent the median and each point is a biological replicate. *$p < 0.05$; Mann–Whitney U test.
