## Supplementary tables for "Ether phospholipids modulate somatosensory responses by tuning multiple receptor functions in *Drosophila*"

Table S1. LC-MS analysis of PE in AGPS KO *Drosophila* whole body[a]

| | Control | *AGPS*$^{KO/+}$ | | *AGPS*$^{KO/KO}$ | |
|---|---|---|---|---|---|
| PE(30:0) | N.D. | N.D. | | N.D. | |
| PE(30:1) | 2.051 ± 0.123 | 1.788 ± 0.071 | | 1.793 ± 0.074 | |
| PE(30:2) | 0.177 ± 0.011 | 0.157 ± 0.003 | | 0.158 ± 0.009 | |
| PE(32:0) | 0.086 ± 0.003 | 0.076 ± 0.006 | | 0.089 ± 0.003 | |
| PE(32:1) | 11.052 ± 0.464 | 9.879 ± 0.159 | | 11.032 ± 0.409 | |
| PE(32:2) | 5.268 ± 0.242 | 4.790 ± 0.198 | | 4.700 ± 0.175 | |
| PE(32:3) | 0.125 ± 0.007 | 0.103 ± 0.005 | | 0.104 ± 0.006 | * |
| PE(34:0) | N.D. | N.D. | | N.D. | |
| PE(34:1) | 34.794 ± 0.709 | 35.427 ± 0.657 | | 34.382 ± 0.847 | |
| PE(34:2) | 19.901 ± 0.691 | 18.671 ± 0.288 | | 19.888 ± 0.793 | |
| PE(34:3) | 3.481 ± 0.100 | 3.031 ± 0.054 | | 3.234 ± 0.078 | ** |
| PE(34:4) | 0.166 ± 0.007 | 0.135 ± 0.005 | | 0.143 ± 0.008 | * |
| PE(36:0) | N.D. | N.D. | | N.D. | |
| PE(36:1) | 4.393 ± 0.766 | 4.492 ± 0.257 | | 5.356 ± 0.638 | |
| PE(36:2) | 13.165 ± 0.581 | 16.157 ± 0.532 | * | 14.640 ± 1.079 | |
| PE(36:3) | 5.035 ± 0.074 | 4.601 ± 0.057 | ** | 4.485 ± 0.097 | ** |
| PE(36:4) | 0.867 ± 0.032 | 0.718 ± 0.009 | | 0.812 ± 0.023 | ** |
| PE(36:5) | 0.059 ± 0.002 | 0.047 ± 0.002 | | 0.054 ± 0.002 | ** |
| PE(36:6) | N.D. | N.D. | | N.D. | |
| PE(38:0) | N.D. | N.D. | | N.D. | |
| PE(38:1) | 0.530 ± 0.036 | 0.575 ± 0.038 | | 0.496 ± 0.036 | |
| PE(38:2) | 0.394 ± 0.067 | 0.508 ± 0.039 | | 0.515 ± 0.010 | |
| PE(38:3) | 0.050 ± 0.008 | 0.068 ± 0.005 | | 0.068 ± 0.007 | |
| PE(38:4) | N.D. | N.D. | | N.D. | |
| PE(38:5) | N.D. | N.D. | | N.D. | |
| PE(38:6) | N.D. | N.D. | | N.D. | |
| PE(40:0) | N.D. | N.D. | | N.D. | |
| PE(40:1) | N.D. | N.D. | | N.D. | |
| PE(40:2) | N.D. | N.D. | | N.D. | |
| PE(40:3) | N.D. | N.D. | | N.D. | |
| PE(40:4) | N.D. | N.D. | | N.D. | |
| PE(40:5) | N.D. | N.D. | | N.D. | |
| PE(40:6) | N.D. | N.D. | | N.D. | |
| PE(40:7) | N.D. | N.D. | | N.D. | |
| PE(40:8) | N.D. | N.D. | | N.D. | |
| ePE(32:0) | N.D. | N.D. | | N.D. | |
| ePE(32:1) | N.D. | N.D. | | N.D. | |
| ePE(34:0) | N.D. | N.D. | | N.D. | |
| ePE(34:1) | 0.102 ± 0.019 | 0.096 ± 0.010 | | N.D. | |
| ePE(34:2) | 0.006 ± 0.001 | 0.006 ± 0.000 | | N.D. | |
| ePE(34:3) | N.D. | N.D. | | N.D. | |
| ePE(36:0) | N.D. | N.D. | | N.D. | |
| ePE(36:1) | 0.114 ± 0.009 | 0.124 ± 0.010 | | N.D. | |
| ePE(36:2) | 0.250 ± 0.045 | 0.274 ± 0.014 | | N.D. | |
| ePE(36:3) | 0.134 ± 0.010 | 0.187 ± 0.008 | ** | N.D. | |
| ePE(36:4) | N.D. | N.D. | | N.D. | |
| ePE(36:5) | N.D. | N.D. | | N.D. | |
| ePE(38:3) | 0.029 ± 0.006 | 0.035 ± 0.003 | | N.D. | |
| ePE(38:4) | N.D. | N.D. | | N.D. | |
| ePE(38:5) | N.D. | N.D. | | N.D. | |
| ePE(38:6) | N.D. | N.D. | | N.D. | |
| Total ePE | 0.636 ± 0.082 | 0.722 ± 0.040 | | N.D. | |

[a]The lipid compositions of PE species (area% of total PE), Mean±SEM,
N.D. represents phospholipid species that were lower than detectable level.
*p<0.05; **p<0.01, Dunnett's test (vs. control)

Table S2. LC-MS analysis of PC in AGPS KO *Drosophila* whole body[a]

| | Control | $AGPS^{KO/+}$ | | $AGPS^{KO/KO}$ | |
|---|---|---|---|---|---|
| PC(26:0) | 0.594 ± 0.019 | 0.576 ± 0.023 | | 0.540 ± 0.021 | |
| PC(26:1) | 0.202 ± 0.011 | 0.191 ± 0.016 | | 0.210 ± 0.006 | |
| PC(28:0) | 0.592 ± 0.018 | 0.600 ± 0.015 | | 0.578 ± 0.016 | |
| PC(28:1) | 1.612 ± 0.041 | 1.591 ± 0.044 | | 1.369 ± 0.036 | ** |
| PC(28:2) | 0.432 ± 0.020 | 0.432 ± 0.011 | | 0.317 ± 0.017 | ** |
| PC(30:0) | 0.451 ± 0.014 | 0.437 ± 0.013 | | 0.444 ± 0.009 | |
| PC(30:1) | 8.085 ± 0.261 | 7.885 ± 0.056 | | 8.081 ± 0.087 | |
| PC(30:2) | 1.749 ± 0.053 | 1.697 ± 0.013 | | 1.520 ± 0.015 | ** |
| PC(32:0) | 0.083 ± 0.013 | 0.099 ± 0.010 | | 0.100 ± 0.011 | |
| PC(32:1) | 16.324 ± 0.380 | 15.735 ± 0.206 | | 16.443 ± 0.162 | |
| PC(32:2) | 16.512 ± 0.460 | 15.738 ± 0.403 | | 16.025 ± 0.117 | |
| PC(32:3) | 0.535 ± 0.012 | 0.494 ± 0.006 | * | 0.500 ± 0.006 | * |
| PC(34:0) | N.D. | N.D. | | N.D. | |
| PC(34:1) | 14.372 ± 1.345 | 15.744 ± 0.438 | | 14.351 ± 0.094 | |
| PC(34:2) | 18.826 ± 0.601 | 19.262 ± 0.320 | | 19.528 ± 0.285 | |
| PC(34:3) | 6.643 ± 0.064 | 6.065 ± 0.120 | | 6.835 ± 0.099 | ** |
| PC(34:4) | 0.286 ± 0.011 | 0.244 ± 0.005 | | 0.286 ± 0.013 | * |
| PC(36:0) | N.D. | N.D. | | N.D. | |
| PC(36:1) | 0.449 ± 0.014 | 0.462 ± 0.029 | | 0.383 ± 0.015 | |
| PC(36:2) | 5.005 ± 0.450 | 5.547 ± 0.194 | | 5.114 ± 0.309 | |
| PC(36:3) | 5.252 ± 0.158 | 5.369 ± 0.108 | | 5.219 ± 0.323 | |
| PC(36:4) | 1.287 ± 0.044 | 1.161 ± 0.031 | | 1.375 ± 0.051 | |
| PC(36:5) | 0.053 ± 0.001 | 0.044 ± 0.005 | | 0.058 ± 0.003 | |
| PC(36:6) | N.D. | N.D. | | N.D. | |
| PC(38:0) | 0.105 ± 0.007 | 0.118 ± 0.005 | | 0.117 ± 0.006 | |
| PC(38:1) | 0.300 ± 0.051 | 0.275 ± 0.004 | | 0.325 ± 0.024 | |
| PC(38:2) | 0.130 ± 0.013 | 0.128 ± 0.007 | | 0.161 ± 0.004 | |
| PC(38:3) | N.D. | N.D. | | N.D. | |
| PC(38:4) | N.D. | N.D. | | N.D. | |
| PC(38:5) | N.D. | N.D. | | N.D. | |
| PC(38:6) | N.D. | N.D. | | N.D. | |
| PC(40:1) | 0.023 ± 0.004 | 0.032 ± 0.001 | | 0.025 ± 0.002 | |
| PC(40:2) | 0.074 ± 0.019 | 0.055 ± 0.005 | | 0.065 ± 0.005 | |
| PC(40:3) | 0.024 ± 0.002 | 0.018 ± 0.002 | | 0.023 ± 0.002 | |
| PC(40:4) | N.D. | N.D. | | N.D. | |
| PC(40:5) | N.D. | N.D. | | N.D. | |
| PC(40:6) | N.D. | N.D. | | N.D. | |
| PC(40:7) | N.D. | N.D. | | N.D. | |
| PC(40:8) | N.D. | N.D. | | N.D. | |
| ePC(32:0) | N.D. | N.D. | | N.D. | |
| ePC(32:1) | N.D. | N.D. | | N.D. | |
| ePC(34:0) | N.D. | N.D. | | N.D. | |
| ePC(34:1) | N.D. | N.D. | | N.D. | |
| ePC(34:2) | N.D. | N.D. | | N.D. | |
| ePC(34:3) | N.D. | N.D. | | N.D. | |
| ePC(36:0) | N.D. | N.D. | | N.D. | |
| ePC(36:1) | N.D. | N.D. | | N.D. | |
| ePC(36:2) | N.D. | N.D. | | N.D. | |
| ePC(36:3) | N.D. | N.D. | | N.D. | |
| ePC(36:4) | N.D. | N.D. | | N.D. | |
| Total ePC | N.D. | N.D. | | N.D. | |

[a]The lipid compositions of PC species (area% of total PC), Mean±SEM,
N.D. represents phospholipid species that were lower than detectable level.
*p<0.05; **p<0.01, Dunnett's test (vs. control)

Table S3. LC-MS analysis of PE in AGPS KO *Drosophila* CNS[a]

| | Control | $AGPS^{KO/KO}$ | |
|---|---|---|---|
| PE(30:0) | 0.137 ± 0.009 | 0.212 ± 0.004 | ** |
| PE(30:1) | 0.845 ± 0.067 | 0.966 ± 0.035 | |
| PE(30:2) | 0.045 ± 0.004 | 0.045 ± 0.003 | |
| PE(32:0) | 0.288 ± 0.016 | 0.560 ± 0.018 | ** |
| PE(32:1) | 4.466 ± 0.329 | 5.593 ± 0.181 | * |
| PE(32:2) | 1.108 ± 0.096 | 1.304 ± 0.043 | |
| PE(32:3) | 0.061 ± 0.005 | 0.092 ± 0.004 | ** |
| PE(34:0) | 0.281 ± 0.074 | 0.607 ± 0.032 | * |
| PE(34:1) | 27.125 ± 0.878 | 30.059 ± 0.268 | * |
| PE(34:2) | 12.200 ± 1.024 | 14.970 ± 0.500 | |
| PE(34:3) | 3.703 ± 0.255 | 5.100 ± 0.138 | ** |
| PE(34:4) | 0.372 ± 0.023 | 0.650 ± 0.019 | *** |
| PE(36:0) | 0.009 ± 0.001 | 0.035 ± 0.003 | ** |
| PE(36:1) | 10.205 ± 1.472 | 10.568 ± 0.279 | |
| PE(36:2) | 14.356 ± 0.192 | 16.047 ± 0.322 | ** |
| PE(36:3) | 6.536 ± 0.355 | 10.227 ± 0.048 | ** |
| PE(36:4) | 2.009 ± 0.174 | 3.159 ± 0.056 | ** |
| PE(36:5) | 0.303 ± 0.021 | 0.476 ± 0.011 | ** |
| PE(36:6) | 0.052 ± 0.002 | 0.079 ± 0.005 | ** |
| PE(38:0) | N.D. | N.D. | |
| PE(38:1) | 0.205 ± 0.007 | 0.278 ± 0.009 | *** |
| PE(38:2) | 0.124 ± 0.022 | 0.131 ± 0.007 | |
| PE(38:3) | 0.043 ± 0.004 | 0.065 ± 0.001 | ** |
| PE(38:4) | N.D. | N.D. | |
| PE(38:5) | N.D. | N.D. | |
| PE(38:6) | N.D. | N.D. | |
| PE(40:0) | N.D. | N.D. | |
| PE(40:1) | N.D. | N.D. | |
| PE(40:2) | N.D. | N.D. | |
| PE(40:3) | N.D. | N.D. | |
| PE(40:4) | N.D. | N.D. | |
| PE(40:5) | N.D. | N.D. | |
| PE(40:6) | N.D. | N.D. | |
| PE(40:7) | N.D. | N.D. | |
| PE(40:8) | N.D. | N.D. | |
| ePE(32:0) | 0.021 ± 0.003 | N.D. | |
| ePE(32:1) | 0.041 ± 0.002 | N.D. | |
| ePE(34:0) | 0.101 ± 0.019 | N.D. | |
| ePE(34:1) | 2.142 ± 0.265 | N.D. | |
| ePE(34:2) | 0.095 ± 0.003 | N.D. | |
| ePE(34:3) | 0.054 ± 0.004 | N.D. | |
| ePE(36:0) | 0.018 ± 0.002 | N.D. | |
| ePE(36:1) | 3.721 ± 0.382 | N.D. | |
| ePE(36:2) | 6.159 ± 0.669 | N.D. | |
| ePE(36:3) | 3.665 ± 0.100 | N.D. | |
| ePE(36:4) | N.D. | N.D. | |
| ePE(36:5) | N.D. | N.D. | |
| ePE(38:3) | 0.534 ± 0.106 | N.D. | |
| ePE(38:4) | N.D. | N.D. | |
| ePE(38:5) | N.D. | N.D. | |
| ePE(38:6) | N.D. | N.D. | |
| Total ePE | 16.552 ± 1.473 | N.D. | |

[a]The lipid compositions of PE species (area% of total PE), Mean±SEM,
N.D. represents phospholipid species that were lower than detectable level.
$*p<0.05$; $**p<0.01$; $***p<0.001$, Student's t-test

Table S4. LC-MS analysis of PC in AGPS KO *Drosophila* CNS[a]

| | Control | *AGPS*$^{KO/KO}$ | |
|---|---|---|---|
| PC(26:0) | N.D. | N.D. | |
| PC(26:1) | N.D. | N.D. | |
| PC(28:0) | 0.079 ± 0.005 | 0.077 ± 0.002 | |
| PC(28:1) | 0.049 ± 0.004 | 0.045 ± 0.005 | |
| PC(28:2) | N.D. | N.D. | |
| PC(30:0) | 0.515 ± 0.016 | 0.646 ± 0.019 | ** |
| PC(30:1) | 1.460 ± 0.066 | 1.322 ± 0.041 | |
| PC(30:2) | 0.093 ± 0.005 | 0.075 ± 0.008 | |
| PC(32:0) | 0.756 ± 0.038 | 1.034 ± 0.054 | ** |
| PC(32:1) | 14.705 ± 0.617 | 15.994 ± 0.270 | |
| PC(32:2) | 7.240 ± 0.224 | 7.636 ± 0.242 | |
| PC(32:3) | 0.194 ± 0.008 | 0.205 ± 0.007 | |
| PC(34:0) | 0.098 ± 0.029 | 0.199 ± 0.017 | * |
| PC(34:1) | 18.420 ± 1.625 | 15.634 ± 0.311 | |
| PC(34:2) | 22.119 ± 1.132 | 21.797 ± 0.338 | |
| PC(34:3) | 8.211 ± 0.237 | 9.674 ± 0.190 | ** |
| PC(34:4) | 0.946 ± 0.025 | 1.186 ± 0.027 | *** |
| PC(36:0) | N.D. | N.D. | |
| PC(36:1) | 0.902 ± 0.047 | 1.000 ± 0.030 | |
| PC(36:2) | 8.191 ± 0.898 | 5.770 ± 0.175 | * |
| PC(36:3) | 11.462 ± 0.267 | 11.914 ± 0.262 | |
| PC(36:4) | 3.136 ± 0.107 | 3.903 ± 0.121 | ** |
| PC(36:5) | 0.346 ± 0.011 | 0.474 ± 0.023 | ** |
| PC(36:6) | 0.022 ± 0.002 | 0.034 ± 0.003 | * |
| PC(38:0) | 0.209 ± 0.014 | 0.233 ± 0.012 | |
| PC(38:1) | 0.428 ± 0.077 | 0.568 ± 0.031 | |
| PC(38:2) | 0.121 ± 0.014 | 0.195 ± 0.013 | ** |
| PC(38:3) | N.D. | N.D. | |
| PC(38:4) | N.D. | N.D. | |
| PC(38:5) | N.D. | N.D. | |
| PC(38:6) | N.D. | N.D. | |
| PC(40:1) | 0.066 ± 0.007 | 0.059 ± 0.004 | |
| PC(40:2) | 0.200 ± 0.041 | 0.265 ± 0.024 | |
| PC(40:3) | 0.035 ± 0.004 | 0.061 ± 0.007 | * |
| PC(40:4) | N.D. | N.D. | |
| PC(40:5) | N.D. | N.D. | |
| PC(40:6) | N.D. | N.D. | |
| PC(40:7) | N.D. | N.D. | |
| PC(40:8) | N.D. | N.D. | |
| ePC(32:0) | N.D. | N.D. | |
| ePC(32:1) | N.D. | N.D. | |
| ePC(34:0) | N.D. | N.D. | |
| ePC(34:1) | N.D. | N.D. | |
| ePC(34:2) | N.D. | N.D. | |
| ePC(34:3) | N.D. | N.D. | |
| ePC(36:0) | N.D. | N.D. | |
| ePC(36:1) | N.D. | N.D. | |
| ePC(36:2) | N.D. | N.D. | |
| ePC(36:3) | N.D. | N.D. | |
| ePC(36:4) | N.D. | N.D. | |
| Total ePC | N.D. | N.D. | |

[a]The lipid compositions of PC species (area% of total PC), Mean±SEM,
N.D. represents phospholipid species that were lower than detectable level.
*p<0.05; **p<0.01; ***p<0.001, Student's t-test

Table S5  LC-MS analysis of PE in *Drosophila* CNS of glial or neural AGPS RNAi[a]

| | + > *AGPS* RNAi | | | | repo > *AGPS* RNAi | | | | elav > *AGPS* RNAi | | | |
|---|---|---|---|---|---|---|---|---|---|---|---|---|
| PE(30:0) | 0.267 | ± | 0.008 | | 0.218 | ± | 0.015 | * | 0.258 | ± | 0.008 | |
| PE(30:1) | 0.962 | ± | 0.067 | | 0.822 | ± | 0.026 | | 0.854 | ± | 0.045 | |
| PE(30:2) | 0.040 | ± | 0.006 | | 0.029 | ± | 0.002 | | 0.030 | ± | 0.002 | |
| PE(32:0) | 0.491 | ± | 0.028 | | 0.446 | ± | 0.018 | | 0.604 | ± | 0.036 | * |
| PE(32:1) | 4.871 | ± | 0.216 | | 4.847 | ± | 0.095 | | 5.076 | ± | 0.265 | |
| PE(32:2) | 1.626 | ± | 0.110 | | 1.888 | ± | 0.058 | | 1.597 | ± | 0.057 | |
| PE(32:3) | 0.131 | ± | 0.017 | | 0.125 | ± | 0.010 | | 0.132 | ± | 0.013 | |
| PE(34:0) | 0.887 | ± | 0.025 | | 0.850 | ± | 0.017 | | 1.269 | ± | 0.011 | *** |
| PE(34:1) | 18.215 | ± | 1.047 | | 18.686 | ± | 0.751 | | 17.201 | ± | 0.917 | |
| PE(34:2) | 12.153 | ± | 0.849 | | 11.766 | ± | 0.442 | | 13.626 | ± | 0.969 | |
| PE(34:3) | 7.438 | ± | 0.153 | | 7.450 | ± | 0.228 | | 7.921 | ± | 0.327 | |
| PE(34:4) | 0.612 | ± | 0.055 | | 0.776 | ± | 0.040 | | 0.766 | ± | 0.039 | |
| PE(36:0) | 0.020 | ± | 0.001 | | 0.020 | ± | 0.002 | | 0.031 | ± | 0.001 | *** |
| PE(36:1) | 9.251 | ± | 0.269 | | 8.741 | ± | 0.331 | | 9.825 | ± | 0.235 | |
| PE(36:2) | 10.830 | ± | 0.240 | | 10.882 | ± | 0.060 | | 11.136 | ± | 0.348 | |
| PE(36:3) | 7.913 | ± | 0.549 | | 7.953 | ± | 0.286 | | 8.141 | ± | 0.445 | |
| PE(36:4) | 2.871 | ± | 0.249 | | 2.620 | ± | 0.149 | | 3.064 | ± | 0.279 | |
| PE(36:5) | 0.565 | ± | 0.036 | | 0.502 | ± | 0.023 | | 0.682 | ± | 0.032 | * |
| PE(36:6) | 0.045 | ± | 0.003 | | 0.042 | ± | 0.003 | | 0.055 | ± | 0.005 | |
| PE(38:0) | | N.D. | | | | N.D. | | | | N.D. | | |
| PE(38:1) | 0.340 | ± | 0.020 | | 0.312 | ± | 0.009 | | 0.328 | ± | 0.011 | |
| PE(38:2) | 0.113 | ± | 0.007 | | 0.112 | ± | 0.009 | | 0.124 | ± | 0.004 | |
| PE(38:3) | 0.085 | ± | 0.005 | | 0.081 | ± | 0.004 | | 0.095 | ± | 0.004 | |
| PE(38:4) | | N.D. | | | | N.D. | | | | N.D. | | |
| PE(38:5) | | N.D. | | | | N.D. | | | | N.D. | | |
| PE(38:6) | | N.D. | | | | N.D. | | | | N.D. | | |
| PE(40:0) | | N.D. | | | | N.D. | | | | N.D. | | |
| PE(40:1) | | N.D. | | | | N.D. | | | | N.D. | | |
| PE(40:2) | | N.D. | | | | N.D. | | | | N.D. | | |
| PE(40:3) | | N.D. | | | | N.D. | | | | N.D. | | |
| PE(40:4) | | N.D. | | | | N.D. | | | | N.D. | | |
| PE(40:5) | | N.D. | | | | N.D. | | | | N.D. | | |
| PE(40:6) | | N.D. | | | | N.D. | | | | N.D. | | |
| PE(40:7) | | N.D. | | | | N.D. | | | | N.D. | | |
| PE(40:8) | | N.D. | | | | N.D. | | | | N.D. | | |
| ePE(32:0) | 0.086 | ± | 0.001 | | 0.085 | ± | 0.005 | | 0.073 | ± | 0.004 | |
| ePE(32:1) | 0.107 | ± | 0.002 | | 0.151 | ± | 0.006 | *** | 0.080 | ± | 0.005 | *** |
| ePE(34:0) | 0.204 | ± | 0.007 | | 0.210 | ± | 0.015 | | 0.221 | ± | 0.013 | |
| ePE(34:1) | 3.184 | ± | 0.188 | | 3.738 | ± | 0.269 | | 3.107 | ± | 0.132 | |
| ePE(34:2) | 0.262 | ± | 0.007 | | 0.297 | ± | 0.010 | * | 0.199 | ± | 0.006 | *** |
| ePE(34:3) | | N.D. | | | | N.D. | | | | N.D. | | |
| ePE(36:0) | 0.109 | ± | 0.004 | | 0.096 | ± | 0.003 | | 0.116 | ± | 0.004 | |
| ePE(36:1) | 4.388 | ± | 0.271 | | 4.701 | ± | 0.227 | | 3.218 | ± | 0.140 | ** |
| ePE(36:2) | 6.377 | ± | 0.328 | | 6.490 | ± | 0.498 | | 5.707 | ± | 0.247 | |
| ePE(36:3) | 4.808 | ± | 0.186 | | 4.371 | ± | 0.169 | | 3.996 | ± | 0.081 | ** |
| ePE(36:4) | | N.D. | | | | N.D. | | | | N.D. | | |
| ePE(36:5) | | N.D. | | | | N.D. | | | | N.D. | | |
| ePE(38:3) | 0.749 | ± | 0.036 | | 0.693 | ± | 0.048 | | 0.467 | ± | 0.017 | *** |
| ePE(38:4) | | N.D. | | | | N.D. | | | | N.D. | | |
| ePE(38:5) | | N.D. | | | | N.D. | | | | N.D. | | |
| ePE(38:6) | | N.D. | | | | N.D. | | | | N.D. | | |
| Total ePE | 19.8768 | ± | 0.452 | | 20.386 | ± | 0.670 | | 16.809 | ± | 0.401 | ** |

[a]The  lipid compositions of PE species (area% of total PE), Mean±SEM,
N.D. represents phospholipid species that were lower than detectable level.
*p<0.05; **p<0.01; ***p<0.001, Dunnett's test (vs. control)

Table S6. LC-MS analysis of PC in *Drosophila* CNS of glial or neural AGPS RNAi[a]

| | + > AGPS RNAi | repo > AGPS RNAi | elav > AGPS RNAi |
|---|---|---|---|
| PC(26:0) | N.D. | N.D. | N.D. |
| PC(26:1) | N.D. | N.D. | N.D. |
| PC(28:0) | 0.100 ± 0.016 | 0.080 ± 0.011 | 0.071 ± 0.008 |
| PC(28:1) | 0.100 ± 0.011 | 0.043 ± 0.010 ** | 0.059 ± 0.009 * |
| PC(28:2) | N.D. | N.D. | N.D. |
| PC(30:0) | 0.908 ± 0.011 | 0.692 ± 0.023 *** | 0.815 ± 0.026 * |
| PC(30:1) | 1.894 ± 0.193 | 1.493 ± 0.086 | 1.466 ± 0.133 |
| PC(30:2) | 0.147 ± 0.013 | 0.108 ± 0.013 | 0.091 ± 0.024 |
| PC(32:0) | 1.293 ± 0.148 | 1.096 ± 0.051 | 1.433 ± 0.120 |
| PC(32:1) | 17.701 ± 1.582 | 17.087 ± 0.781 | 16.768 ± 0.992 |
| PC(32:2) | 7.498 ± 0.493 | 8.139 ± 0.115 | 7.382 ± 0.050 |
| PC(32:3) | 0.432 ± 0.012 | 0.334 ± 0.029 | 0.370 ± 0.035 |
| PC(34:0) | 0.356 ± 0.022 | 0.318 ± 0.011 | 0.499 ± 0.010 *** |
| PC(34:1) | 14.370 ± 0.441 | 14.909 ± 0.450 | 12.753 ± 0.628 |
| PC(34:2) | 22.712 ± 0.244 | 22.967 ± 0.353 | 23.392 ± 0.398 |
| PC(34:3) | 9.108 ± 0.772 | 9.358 ± 0.297 | 10.491 ± 0.440 |
| PC(34:4) | 1.211 ± 0.035 | 1.263 ± 0.031 | 1.533 ± 0.064 *** |
| PC(36:0) | N.D. | N.D. | N.D. |
| PC(36:1) | 0.962 ± 0.038 | 0.872 ± 0.045 | 0.878 ± 0.025 |
| PC(36:2) | 7.781 ± 0.332 | 7.677 ± 0.180 | 6.890 ± 0.173 * |
| PC(36:3) | 7.959 ± 0.320 | 8.444 ± 0.314 | 8.521 ± 0.210 |
| PC(36:4) | 3.986 ± 0.175 | 3.671 ± 0.099 | 4.910 ± 0.096 ** |
| PC(36:5) | 0.424 ± 0.032 | 0.331 ± 0.011 * | 0.562 ± 0.021 ** |
| PC(36:6) | 0.014 ± 0.003 | 0.012 ± 0.001 | 0.019 ± 0.004 |
| PC(38:0) | 0.208 ± 0.014 | 0.215 ± 0.008 | 0.182 ± 0.013 |
| PC(38:1) | 0.373 ± 0.009 | 0.379 ± 0.042 | 0.379 ± 0.043 |
| PC(38:2) | 0.092 ± 0.005 | 0.117 ± 0.009 | 0.120 ± 0.011 |
| PC(38:3) | N.D. | N.D. | N.D. |
| PC(38:4) | N.D. | N.D. | N.D. |
| PC(38:5) | N.D. | N.D. | N.D. |
| PC(38:6) | N.D. | N.D. | N.D. |
| PC(40:1) | 0.154 ± 0.005 | 0.182 ± 0.016 | 0.165 ± 0.023 |
| PC(40:2) | 0.189 ± 0.008 | 0.180 ± 0.015 | 0.207 ± 0.020 |
| PC(40:3) | 0.029 ± 0.002 | 0.033 ± 0.005 | 0.045 ± 0.005 * |
| PC(40:4) | N.D. | N.D. | N.D. |
| PC(40:5) | N.D. | N.D. | N.D. |
| PC(40:6) | N.D. | N.D. | N.D. |
| PC(40:7) | N.D. | N.D. | N.D. |
| PC(40:8) | N.D. | N.D. | N.D. |
| ePC(32:0) | N.D. | N.D. | N.D. |
| ePC(32:1) | N.D. | N.D. | N.D. |
| ePC(34:0) | N.D. | N.D. | N.D. |
| ePC(34:1) | N.D. | N.D. | N.D. |
| ePC(34:2) | N.D. | N.D. | N.D. |
| ePC(34:3) | N.D. | N.D. | N.D. |
| ePC(36:0) | N.D. | N.D. | N.D. |
| ePC(36:1) | N.D. | N.D. | N.D. |
| ePC(36:2) | N.D. | N.D. | N.D. |
| ePC(36:3) | N.D. | N.D. | N.D. |
| ePC(36:4) | N.D. | N.D. | N.D. |
| Total ePC | N.D. | N.D. | N.D. |

[a]The lipid compositions of PC species (area% of total PC), Mean±SEM,
N.D. represents phospholipid species that were lower than detectable level.
*p<0.05; **p<0.01; ***p<0.001, Dunnett’s test (vs. control)

Table S7. LC-MS analysis of PE in ePL producing S2R+ cells[a]

| | Control/S2R+ | 16-AG/S2R+ | | Control/S2R+ dAGPS | 18-OH/S2R+ dAGPS | |
|---|---|---|---|---|---|---|
| PE(30:0) | 0.129 ± 0.009 | 0.264 ± 0.006 | *** | 0.095 ± 0.004 | 0.098 ± 0.006 | |
| PE(30:1) | 3.396 ± 0.165 | 2.515 ± 0.025 | * | 3.565 ± 0.073 | 3.447 ± 0.066 | |
| PE(30:2) | N.D. | N.D. | | N.D. | N.D. | |
| PE(32:0) | 0.108 ± 0.007 | 0.380 ± 0.016 | *** | 0.075 ± 0.007 | 0.074 ± 0.007 | |
| PE(32:1) | 10.057 ± 0.323 | 7.758 ± 0.085 | ** | 11.119 ± 0.112 | 9.804 ± 0.111 | *** |
| PE(32:2) | 3.611 ± 0.046 | 4.096 ± 0.109 | * | 3.575 ± 0.066 | 3.697 ± 0.119 | |
| PE(32:3) | N.D. | N.D. | | N.D. | N.D. | |
| PE(34:0) | N.D. | N.D. | | N.D. | N.D. | |
| PE(34:1) | 20.306 ± 0.402 | 16.968 ± 0.290 | *** | 26.409 ± 0.348 | 22.000 ± 0.227 | *** |
| PE(34:2) | 15.988 ± 0.472 | 13.619 ± 0.293 | ** | 14.009 ± 0.177 | 14.451 ± 0.084 | |
| PE(34:3) | 0.412 ± 0.019 | 0.363 ± 0.014 | | 0.368 ± 0.017 | 0.330 ± 0.010 | |
| PE(34:4) | N.D. | N.D. | | N.D. | N.D. | |
| PE(36:0) | N.D. | N.D. | | N.D. | N.D. | |
| PE(36:1) | 7.986 ± 0.136 | 6.854 ± 0.167 | ** | 6.169 ± 0.194 | 5.298 ± 0.152 | * |
| PE(36:2) | 28.911 ± 0.670 | 26.404 ± 0.547 | * | 27.073 ± 0.222 | 28.302 ± 0.252 | * |
| PE(36:3) | 1.521 ± 0.066 | 1.197 ± 0.061 | * | 1.107 ± 0.034 | 1.156 ± 0.037 | |
| PE(36:4) | 0.826 ± 0.023 | 0.754 ± 0.015 | * | 0.640 ± 0.015 | 0.562 ± 0.017 | * |
| PE(36:5) | 0.632 ± 0.024 | 0.676 ± 0.027 | | 0.599 ± 0.024 | 0.547 ± 0.018 | |
| PE(36:6) | 0.134 ± 0.008 | 0.175 ± 0.011 | * | 0.188 ± 0.007 | 0.169 ± 0.002 | |
| PE(38:0) | N.D. | N.D. | | N.D. | N.D. | |
| PE(38:1) | 2.503 ± 0.042 | 2.061 ± 0.067 | ** | 2.382 ± 0.094 | 2.260 ± 0.105 | |
| PE(38:2) | 1.413 ± 0.040 | 1.261 ± 0.048 | * | 0.950 ± 0.025 | 0.813 ± 0.014 | ** |
| PE(38:3) | N.D. | N.D. | | N.D. | N.D. | |
| PE(38:4) | 0.473 ± 0.015 | 0.512 ± 0.008 | | 0.390 ± 0.007 | 0.357 ± 0.016 | |
| PE(38:5) | 1.112 ± 0.034 | 0.955 ± 0.092 | | 0.870 ± 0.046 | 0.852 ± 0.078 | |
| PE(38:6) | 0.465 ± 0.028 | 0.413 ± 0.019 | | 0.398 ± 0.009 | 0.381 ± 0.010 | |
| PE(40:0) | N.D. | N.D. | | N.D. | N.D. | |
| PE(40:1) | N.D. | N.D. | | N.D. | N.D. | |
| PE(40:2) | N.D. | N.D. | | N.D. | N.D. | |
| PE(40:3) | N.D. | N.D. | | N.D. | N.D. | |
| PE(40:4) | N.D. | N.D. | | N.D. | N.D. | |
| PE(40:5) | N.D. | N.D. | | N.D. | N.D. | |
| PE(40:6) | N.D. | N.D. | | N.D. | N.D. | |
| PE(40:7) | N.D. | N.D. | | N.D. | N.D. | |
| PE(40:8) | N.D. | N.D. | | N.D. | N.D. | |
| ePE(32:0) | N.D. | 0.220 ± 0.011 | | N.D. | N.D. | |
| ePE(32:1) | N.D. | 7.248 ± 0.119 | | N.D. | 0.136 ± 0.007 | |
| ePE(34:0) | N.D. | N.D. | | N.D. | N.D. | |
| ePE(34:1) | N.D. | 4.515 ± 0.133 | | N.D. | 2.612 ± 0.087 | |
| ePE(34:2) | N.D. | 0.358 ± 0.005 | | N.D. | N.D. | |
| ePE(34:3) | N.D. | N.D. | | N.D. | N.D. | |
| ePE(34:4) | N.D. | N.D. | | N.D. | N.D. | |
| ePE(36:0) | N.D. | N.D. | | N.D. | N.D. | |
| ePE(36:1) | 0.014 ± 0.003 | 0.101 ± 0.003 | *** | 0.017 ± 0.003 | 2.657 ± 0.114 | *** |
| ePE(36:2) | N.D. | N.D. | | N.D. | N.D. | |
| ePE(36:3) | N.D. | N.D. | | N.D. | N.D. | |
| ePE(36:4) | N.D. | 0.171 ± 0.005 | | N.D. | N.D. | |
| ePE(36:5) | N.D. | 0.161 ± 0.005 | | N.D. | N.D. | |
| ePE(38:0) | N.D. | N.D. | | N.D. | N.D. | |
| ePE(38:1) | N.D. | N.D. | | N.D. | N.D. | |
| ePE(38:2) | N.D. | N.D. | | N.D. | N.D. | |
| ePE(38:3) | N.D. | N.D. | | N.D. | N.D. | |
| ePE(38:4) | N.D. | N.D. | | N.D. | N.D. | |
| ePE(38:5) | N.D. | N.D. | | N.D. | N.D. | |
| ePE Total | 0.014 ± 0.003 | 12.774 ± 0.254 | *** | 0.017 ± 0.003 | 5.405 ± 0.182 | *** |

[a]The lipid compositions of PE species (area% of total PE), Mean±SEM, *p<0.05; **p<0.01; ***p<0.001, Student's t-test
N.D. represents phospholipid species that were lower than detectable level.

Table S8. LC-MS analysis of PC in ePL producing S2R+ cells[a]

| | Control/S2R+ | 16-AG/S2R+ | | Control/S2R+ dAGPS | 18-OH/S2R+ dAGPS | |
|---|---|---|---|---|---|---|
| PC(28:0) | 0.221 ± 0.008 | 0.247 ± 0.012 | | 0.496 ± 0.012 | 0.463 ± 0.026 | |
| PC(28:1) | 0.159 ± 0.006 | 0.246 ± 0.008 | *** | 0.322 ± 0.011 | 0.313 ± 0.009 | |
| PC(30:0) | 1.229 ± 0.055 | 1.017 ± 0.051 | * | 2.159 ± 0.056 | 2.069 ± 0.035 | |
| PC(30:1) | 2.775 ± 0.036 | 3.029 ± 0.047 | ** | 3.958 ± 0.080 | 3.694 ± 0.066 | |
| PC(30:2) | 0.525 ± 0.027 | 0.608 ± 0.010 | * | 0.580 ± 0.019 | 0.561 ± 0.020 | |
| PC(32:0) | 1.163 ± 0.049 | 0.631 ± 0.046 | *** | 1.296 ± 0.034 | 1.236 ± 0.064 | |
| PC(32:1) | 13.258 ± 0.111 | 14.783 ± 0.441 | * | 17.177 ± 0.328 | 16.861 ± 0.243 | |
| PC(32:2) | 10.957 ± 0.252 | 13.341 ± 0.120 | *** | 11.205 ± 0.148 | 10.217 ± 0.207 | * |
| PC(32:3) | 0.159 ± 0.006 | 0.141 ± 0.009 | | 0.120 ± 0.003 | 0.114 ± 0.008 | |
| PC(34:0) | N.D. | N.D. | | N.D. | N.D. | |
| PC(34:1) | 15.063 ± 0.309 | 14.524 ± 0.459 | | 18.779 ± 0.246 | 19.891 ± 0.426 | |
| PC(34:2) | 25.640 ± 0.509 | 23.804 ± 0.390 | * | 21.409 ± 0.225 | 20.647 ± 0.176 | |
| PC(34:3) | 0.718 ± 0.026 | 0.758 ± 0.024 | | 0.574 ± 0.027 | 0.548 ± 0.010 | |
| PC(34:4) | 0.407 ± 0.010 | 0.412 ± 0.005 | | 0.358 ± 0.022 | 0.333 ± 0.003 | |
| PC(36:0) | N.D. | N.D. | | N.D. | N.D. | |
| PC(36:1) | 1.139 ± 0.042 | 0.990 ± 0.031 | * | 0.737 ± 0.024 | 0.831 ± 0.036 | |
| PC(36:2) | 16.726 ± 0.157 | 15.117 ± 0.268 | ** | 13.830 ± 0.095 | 15.165 ± 0.135 | |
| PC(36:3) | 1.435 ± 0.064 | 1.304 ± 0.067 | | 0.955 ± 0.046 | 0.957 ± 0.056 | |
| PC(36:4) | 1.213 ± 0.008 | 1.273 ± 0.033 | | 0.929 ± 0.035 | 0.955 ± 0.029 | |
| PC(36:5) | 1.447 ± 0.043 | 1.616 ± 0.033 | * | 1.210 ± 0.019 | 1.156 ± 0.034 | |
| PC(36:6) | 0.736 ± 0.040 | 0.658 ± 0.078 | | 0.462 ± 0.021 | 0.455 ± 0.034 | |
| PC(38:0) | N.D. | N.D. | | N.D. | N.D. | |
| PC(38:1) | N.D. | N.D. | | N.D. | N.D. | |
| PC(38:2) | N.D. | N.D. | | N.D. | N.D. | |
| PC(38:3) | N.D. | N.D. | | N.D. | N.D. | |
| PC(38:4) | 0.734 ± 0.035 | 0.837 ± 0.036 | | 0.564 ± 0.025 | 0.528 ± 0.007 | ** |
| PC(38:5) | 2.011 ± 0.086 | 1.790 ± 0.147 | | 1.139 ± 0.086 | 1.220 ± 0.123 | |
| PC(38:6) | 1.357 ± 0.030 | 1.166 ± 0.026 | ** | 1.035 ± 0.013 | 1.041 ± 0.032 | |
| PC(40:1) | N.D. | N.D. | | N.D. | N.D. | |
| PC(40:2) | N.D. | N.D. | | N.D. | N.D. | |
| PC(40:3) | N.D. | N.D. | | N.D. | N.D. | |
| PC(40:4) | N.D. | N.D. | | N.D. | N.D. | |
| PC(40:5) | 0.128 ± 0.010 | 0.138 ± 0.008 | | 0.119 ± 0.015 | 0.122 ± 0.005 | |
| PC(40:6) | 0.293 ± 0.013 | 0.248 ± 0.007 | * | 0.248 ± 0.027 | 0.246 ± 0.021 | |
| PC(40:7) | 0.483 ± 0.021 | 0.308 ± 0.034 | ** | 0.325 ± 0.026 | 0.327 ± 0.042 | |
| PC(40:8) | N.D. | N.D. | | N.D. | N.D. | |
| ePC(30:0) | N.D. | N.D. | | N.D. | N.D. | |
| ePC(30:1) | N.D. | 0.053 ± 0.002 | | N.D. | N.D. | |
| ePC(30:2) | N.D. | N.D. | | N.D. | N.D. | |
| ePC(32:0) | N.D. | N.D. | | N.D. | N.D. | |
| ePC(32:1) | N.D. | 0.715 ± 0.021 | | N.D. | N.D. | |
| ePC(32:2) | N.D. | N.D. | | N.D. | N.D. | |
| ePC(34:0) | N.D. | N.D. | | N.D. | N.D. | |
| ePC(34:1) | 0.025 ± 0.003 | 0.245 ± 0.005 | *** | 0.012 ± 0.001 | 0.048 ± 0.001 | |
| ePC(34:2) | N.D. | N.D. | | N.D. | N.D. | |
| ePC(34:3) | N.D. | N.D. | | N.D. | N.D. | |
| ePC(36:0) | N.D. | N.D. | | N.D. | N.D. | |
| ePC(36:1) | N.D. | N.D. | | N.D. | N.D. | |
| ePC(36:2) | N.D. | N.D. | | N.D. | N.D. | |
| ePC(36:3) | N.D. | N.D. | | N.D. | N.D. | |
| ePC(36:4) | N.D. | N.D. | | N.D. | N.D. | |
| ePC(38:2) | N.D. | N.D. | | N.D. | N.D. | |
| ePC(38:3) | N.D. | N.D. | | N.D. | N.D. | |
| ePC(38:4) | N.D. | N.D. | | N.D. | N.D. | |
| ePC(38:5) | N.D. | N.D. | | N.D. | N.D. | |
| ePC Total | 0.025 ± 0.003 | 1.012 ± 0.026 | *** | 0.012 ± 0.001 | 0.048 ± 0.001 | |

[a]The lipid compositions of PC species (area% of total PC), Mean±SEM, *p<0.05; **p<0.01; ***p<0.001, Student's t-test
N.D. represents phospholipid species that were lower than detectable level.
